# Cables1 links Slit/Robo and Wnt/Frizzled signaling in commissural axon guidance

**DOI:** 10.1101/2021.10.19.464945

**Authors:** Nikole Zuñiga, Esther T. Stoeckli

## Abstract

During neural circuit formation, axons navigate from one intermediate target to the next, until they find their final target. At intermediate targets, axons switch from being attracted to being repelled by changing the guidance receptors on the growth cone surface. For smooth navigation of the intermediate target and the continuation of their journey, the switch in receptor expression has to be orchestrated in a precisely timed manner. As an alternative to changes in expression, receptor function could be regulated by phosphorylation of receptors or components of signaling pathways. We identifed Cables1 as a linker between floor-plate exit of commissural axons regulated by Slit/Robo signaling and the rostral turn of post-crossing axons regulated by Wnt/Frizzled signaling. Cables1 localizes β-Catenin phosphorylated at tyrosine 489 by Abelson kinase to the distal axon, which in turn is necessary for the correct navigation of post-crossing commissural axons in the developing chicken spinal cord.

## INTRODUCTION

During the establishment of neural circuits, axons need to connect to distant targets. On their journey, axons are directed by guidance cues provided by cells along their trajectory and from intermediate targets cutting down the long travelling distance into shorter segments. However, navigation of intermediate targets requires precise control of expression and signaling of guidance receptors (De Ramon Francàs et al., 2017; Stoeckli, 2018; Chédotal, 2019; Ducuing et al., 2019). For example, axons of the dI1 subpopulation of commissural axons cross the floor plate, the ventral midline of the spinal cord, without delay. To this end, the attractive response to the intermediate target, the floor plate, has to be turned into a repulsive response upon arrival, in order to prevent lingering of axons in the midline area, but also to prevent axon guidance errors, due to premature expression of guidance receptors sensing repulsive cues, which would prevent axons from entering the midline area.

Commissural axons are entering the floor-plate area due to the interaction between Contactin-2 (aka Axonin-1) on axons and NrCAM on floor-plate cells (Stoeckli and Landmesser, 1995; Stoeckli et al., 1997). Upon contact with the floor plate, commissural growth cones start expressing Robo1 in a Calsyntenin1- and RabGDI-dependent manner (Alther et al., 2016). The temporally regulated trafficking of Robo receptors to the growth cone surface allows detection of the repulsive Slits only upon entry into the floor-plate area, preventing erroneous ipsilateral turns of axons. Expression of Robo1 on the growth cone surface thus expels axons from the floor plate (Philipp et al., 2012; Alther et al., 2016; Pignata et al., 2019).

Post-crossing axons express Hhip receptors, induced by Shh binding to Glypican1 (Wilson and Stoeckli, 2013), to respond to a repulsive gradient of Shh with high levels in the caudal floor-plate area (Bourikas et al.,2005). At the same time, Shh also shapes an attractive Wnt gradient along the anteroposterior axis, with higher Wnt activity levels anteriorly (Domanitskaya et al., 2010; Lyuksyutova et al., 2003). Components of both canonical and non-canonical Wnt signaling have been implicated in dI1 post-crossing commissural axon guidance along the longitudinal axis suggesting that this strict separation of Wnt signaling into different pathways is not applicable to Wnts’ role in axon guidance (Lyuksyutova et al., 2003; Avilés and Stoeckli, 2016; van Amerongen, 2012). More recently, this has been confirmed again by a detailed study of Wnt signaling in midline crossing at the chiasm (Morenilla-Palao et al., 2020).

Despite the fact that midline crossing appears to be a rather simple, binary decision – to cross or not to cross, its regulation has been shown to be extremely complex, involving a large number of guidance cues and receptors. Because the temporal expression of these guidance receptors on growth cones has to be tightly regulated in order to ensure smooth navigation of an intermediate target, the mechanisms of receptor expression on the growth cone surface have been of great interest. In addition to the subtype-specific response of axons, as shown recently for the ipsi-versus contralaterally projecting retinal ganglion cells (Morenilla-Palao et al., 2020), signaling pathways need to be temporally regulated in the same type of axons. For example, premature expression of receptors for the morphogens presented as gradients along the longitudinal axis of the spinal cord could induce aberrant axonal decisions to turn rostrally along the ipsi- instead of the contralateral floor-plate border.

Our previous studies demonstrated that the responsiveness of only post-but not pre-crossing commissural axons to the antero-posterior Shh gradient is regulated at the transcriptional level: Hhip expression is triggered by Shh binding to Glypican1 on pre-crossing axons (Wilson and Stoeckli, 2013). Similar to the regulation of Robo1 expression, the surface expression of Fzd3, the Wnt receptors on post-crossing commissural axons, is regulated by specific vesicular trafficking (Alther et al., 2016; Onishi and Zou, 2017).

However, we reasoned that exit from the floor plate and turning into the longitudinal axis along the contralateral floor-plate border might be linked by additional mechanisms. A good candidate for such a linker between Slit/Robo signaling and Wnt signaling was Cables1 (Zukerberg et al., 2000; Rhee et al., 2007). In retinal cells, Cables1 was shown to interact with Abl kinase bound to Robo1 triggered by Slit binding, followed by Cables1-mediated transfer of Abl to β-Catenin. This interaction induced dissociation of β-Catenin from N-Cadherin and allowed for phosphorylation of β-Catenin at tyrosine residue 489 by Abl kinase (Rhee et al., 2007).

Here, we show that in the developing spinal cord Cables is required for midline crossing of commissural axons by linking Slit/Robo signaling to Wnt signaling involving phosphorylation of β-Catenin at tyrosine 489 by Abl kinase.

## RESULTS

### Cables1 is upregulated in dI1 neurons during axonal midline crossing

To study the role of Cables in commissural axon guidance we examined the expression pattern of *Cables1. Cables1* mRNA was broadly detected in the chicken spinal cord at different stages during commissural axon navigation (Figure 1). The *Cables1* gene produces different proteins due to alternative splicing both in human and chicken (Zhang et al., 2005). The probe, we used for in situ hybridization recognizes all isoforms. Using qRT-PCR to distinguish the different isoforms of Cables1 indicated that isoform X1 is the predominant splice variant in the developing spinal cord (Supplementary Figure 1A). The dI1 subpopulation of commissural interneurons showed a peak of *Cables1* expression between stages HH22 and HH24, which correspond to critical points in axon navigation, entry and exit of the floor plate, the intermediate target (Figure 1A,B). At later stages, *Cables1* expression in dI1 commissural neurons was reduced to the same levels as found throughout the spinal cord (Figures 1B). The ubiquitous expression of Cables in the developing spinal cord was confirmed by immunostaining. At HH23, Cables protein was detected throughout the spinal cord, including the dI1 population of commissural neurons, as confirmed by Lhx2 staining (Figure 1C). Importantly, Cables was also found in axons crossing the midline, stained with Contactin2/ Axonin-1. Because the antibody was raised against a domain of Cables1 that is 90% identical to the corresponding region of Cables2, and therefore, the antibody does not distinguish between the two proteins, we also studied the expression of *Cables2*, which shares over all 55% of identity with Cables1 at the protein level. If at all, *Cables2* mRNA was expressed at low levels throughout the neural tube at all stages studied (Supplementary Figure 1A,B).

**Figure 1:**
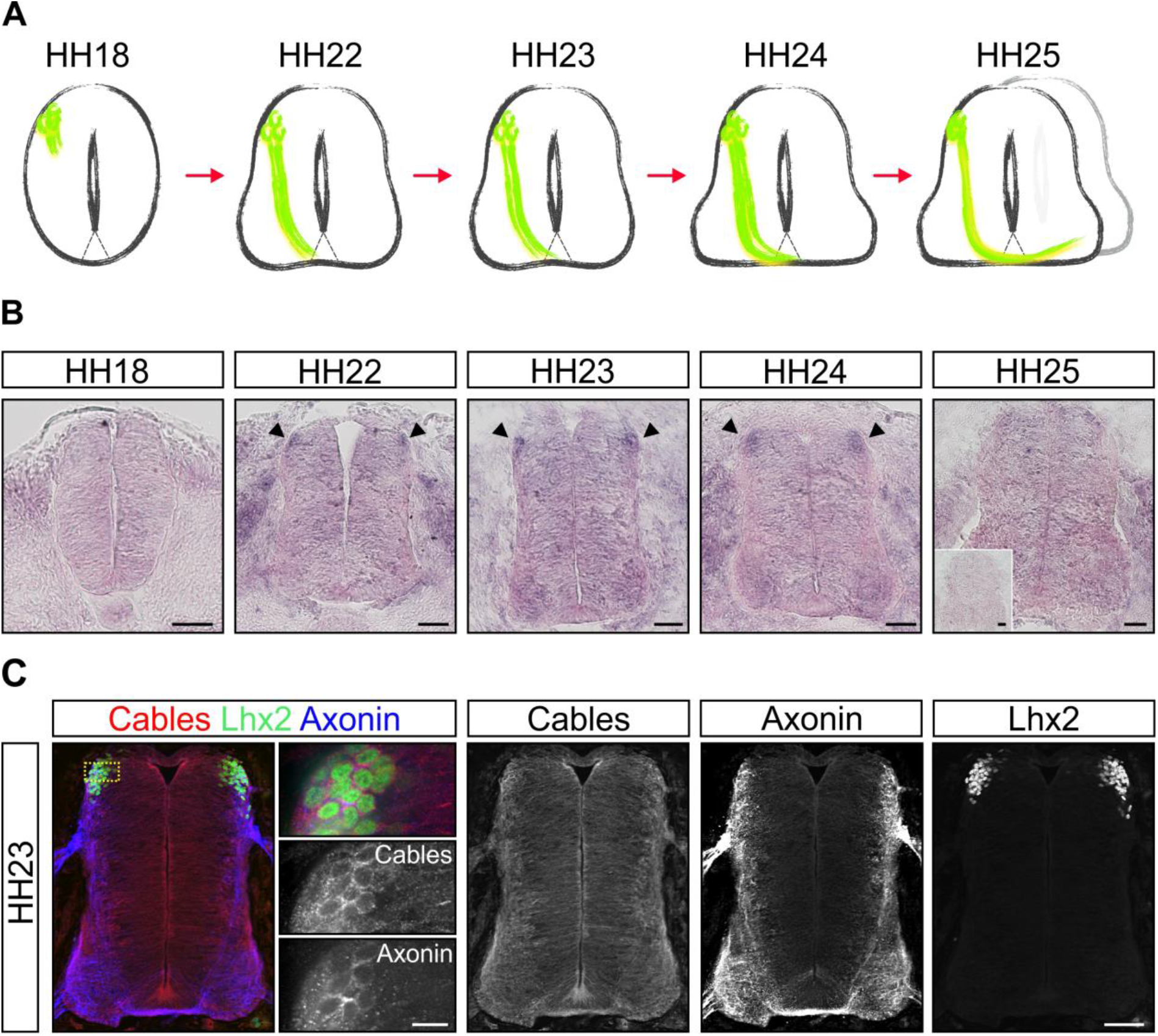
Cables1 is upregulated in dI1 commissural neurons during axonal midline crossing. Expression of Cables1 mRNA is shown in relationship to the temporal development of the dI1 subpopulation of commissural neurons (A). At HH18, dI1 commissural neurons start to extend their axons in the dorsal spinal cord. They reach and enter the floor plate at HH22. At HH24, axons exit the floor plate and turn rostral along the contralateral side of the floor plate. (B) Cables1 mRNA is expressed at low levels throughout the developing neural tube. Higher levels are found in dI1 commissural neurons between HH22 and HH24 (arrowheads), the time window of midline crossing and axonal turning into the longitudinal axis. Expression levels in dI1 neurons decrease after midline navigation at HH25. Insert shows hybridization with sense probe. Immunostaining with an anti-Cables antibody confirms its ubiquitous expression in the developing spinal cord at HH23, when dI1 commissural axons cross the ventral midline (C). Cables is also found in axons crossing the floor plate, labeled with an anti-Axonin1 (Contactin2) antibody. Co-staining with an anti-Lhx2 antibody demonstrates expression of Cables1 protein in dI1 neurons. Bar: 50 µm in B.

### Cables1 is required for axons to exit the floor plate and to turn into the longitudinal axis

The transient upregulation of Cables1 in dI1 neurons during axonal midline crossing suggested a role in commissural axon guidance. In order to test for such a role, we performed *in ovo* RNAi at HH17-18 using long dsRNA to downregulate Cables1. We evaluated the efficiency of downregulation by qRT-PCR and observed a 50% reduction for the transcript levels of *Cables1* isoform1 and less than 40% of the protein left when analyzed by Western blot (Supplementary Figure 2A, B). As we transfected about 50% of the cells in the targeted area with the parameters used in this study, silencing *Cables1* by *in ovo* RNAi virtually removed protein and transcript levels in the transfected cells.

For the analysis of Cables1 function, we analyzed the trajectory of dI1 commissural axons traced by injections of DiI in open-book preparations of spinal cords dissected at HH25-26 (Figure 2A). Axons in untreated and in GFP-expressing control embryos crossed the floor plate and turned rostrally into the longitudinal axis along the contralateral floor-plate border (Figure 2B). Only 24.3±6.1% and 20.7±7.2%, respectively, of the injection sites showed axons with aberrant navigation. In contrast, when Cables1 was downregulated, axons failed to turn rostrally and stalled in the floor plate, or at the exit site, at 68.0±7.4% of the DiI injection sites (Figure 2B). These defects were due to the lack of Cables1, because we could rescue axon guidance by co-expressing mouse *Cables1* cDNA specifically in dI1 neurons using the Math1 enhancer (Figures 2B-2D). Under these conditions, the percentage of DiI injection sites with aberrant axonal trajectories was strongly reduced (35.25±6.5%).

**Figure 2:**
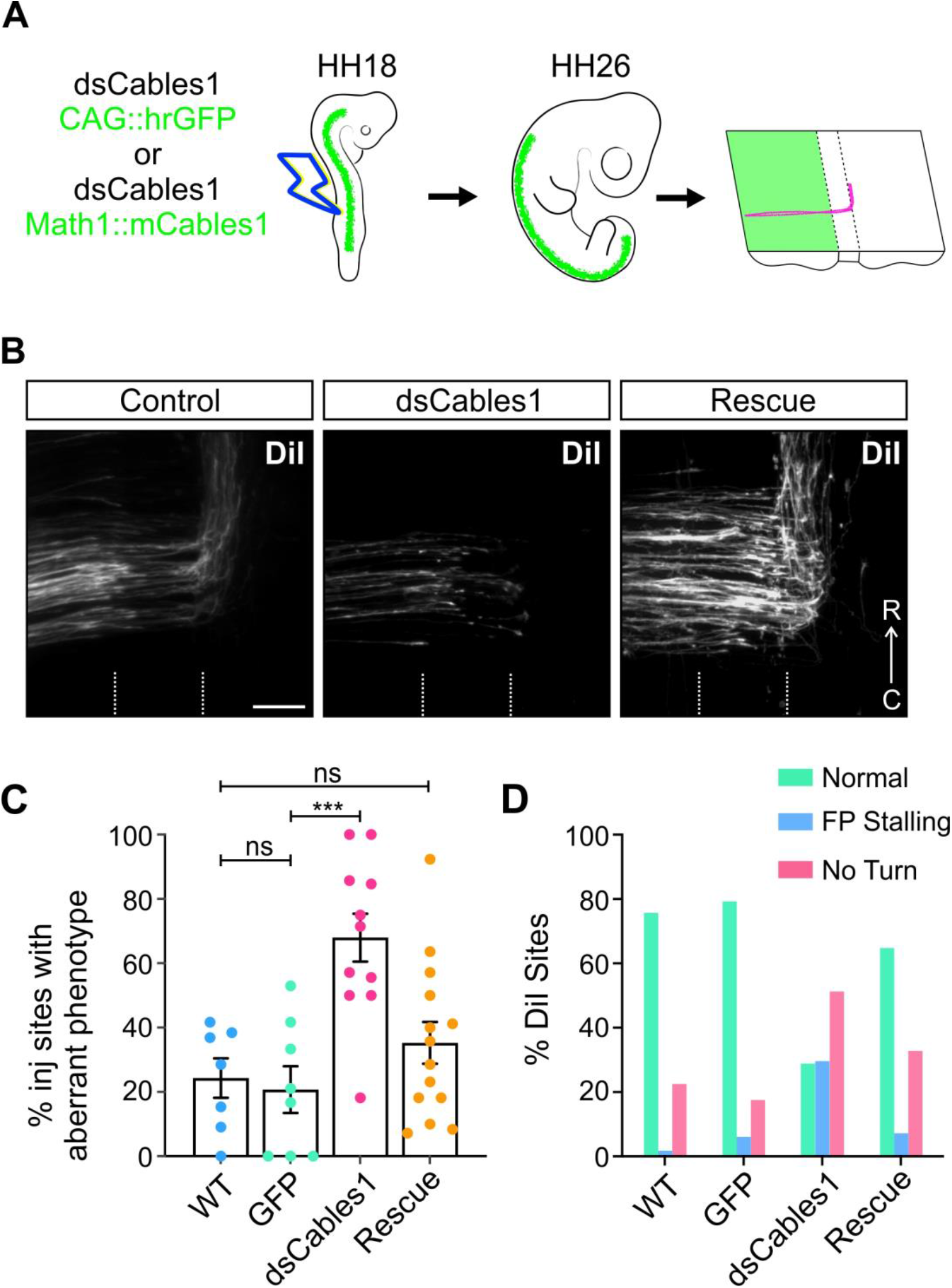
Cables1 is required for axonal navigation at the floor plate. (A) Schematic drawing illustrating loss-of-function experiments using in ovo RNAi. Embryos were injected with dsRNA derived from Cables1 and a plasmid encoding GFP to visualize the transfected area of the neural tube after electroporation of the embryos at HH18. The trajectory of dI1 commissural axons was traced by the lipophilic dye DiI injected into open-book preparations of embryos sacrificed at HH26 (see Experimental Procedures for details). For rescue experiments, embryos were co-injected with dsCables1 and a plasmid encoding mouse Cables1 (mCables1) under the control of the Math1 enhancer for specific expression in dI1 neurons. (B) In control embryos (control), dI1 commissural axons cross the floor plate (indicated by dashed lines) and turn rostral along the contralateral floor-plate border. After silencing Cables1 by the injection and electroporation of dsRNA (dsCables1), most axons failed to cross the floor plate and did not turn into the longitudinal axis. This effect on axonal navigation was specific, because the co-injection and electroporation of dsCables1 together with a plasmid encoding mouse Cables1 that was not targeted by the dsRNA abolished defects in axon guidance (rescue). (C) Quantification of the DiI injection sites with aberrant axonal trajectories resulted in no difference between untreated (control WT; 24.3±6.1%) and GFP-expressing control embryos (GFP; 20.7±7.2%). In contrast, 68.0±7.4% of the injection sites exhibited aberrant axon guidance after silencing Cables1 (***p=0.0004). The effect of dsCables1 on axonal trajectories was abolished when a plasmid encoding mouse Cables1 was co-electroporated (rescue; 35.25±6.5%). This value was not significantly different (ns) from the control groups. Values are mean ± s.e.m. One-way ANOVA with Tukey’s multiple comparisons test. The numbers of injection sites and embryos (in parenthesis) analyzed in each group were: WT n=92 (7 embryos); GFP n=93 (8); dsCables1 n=96 (11); rescue n=160 (14). Bar: 50 µm.

Because we could not exclude that possibility that very low levels of *Cables2* were also expressed throughout the developing spinal cord, including the dI1 neurons, we also silenced *Cables2*. In contrast to our findings for *Cables1*, silencing *Cables2* did not produce any aberrant phenotypes, suggesting that Cables1 function in commissural axons is specific and cannot be compensated by Cables2 (Supplementary Figures 1C,D).

To demonstrate that the phenotype seen after perturbation of Cables1 expression was caused by the lack of axonal expulsion from the floor plate and was not due to a decrease in axonal growth speed, we also analyzed axon guidance phenotypes at HH29-30 (Supplementary Figure 3). At this stage, axons were still stuck in the floor plate or failed to turn into the longitudinal axis after downregulation of Cables1, indicating that the aberrant phenotype was not simply a delay in normal axon growth. We also ruled out an indirect effect on axon guidance by aberrant neuronal differentiation (Supplementary Figure 2C). Taken together, these results suggest that Cables1 is not required for growth of pre-crossing axons, but that it is important for axons to leave the floor plate and turn into the longitudinal axis.

### Cables1 is not required in pre-crossing commissural axons

The absence of an effect of Cables1 on pre-crossing axons was confirmed *in vitro* (Figure 3). We specifically labelled dI1 commissural neurons by electroporation of embryos with a plasmid encoding farnesylated td-Tomato under the control of the Math1 promoter (Figure 3A). When we cultured explants of spinal cords dissected at HH21-22, we found no difference in outgrowth of td-Tomato-positive axons between experimental and control explants (Figure 3B, C).

**Figure 3:**
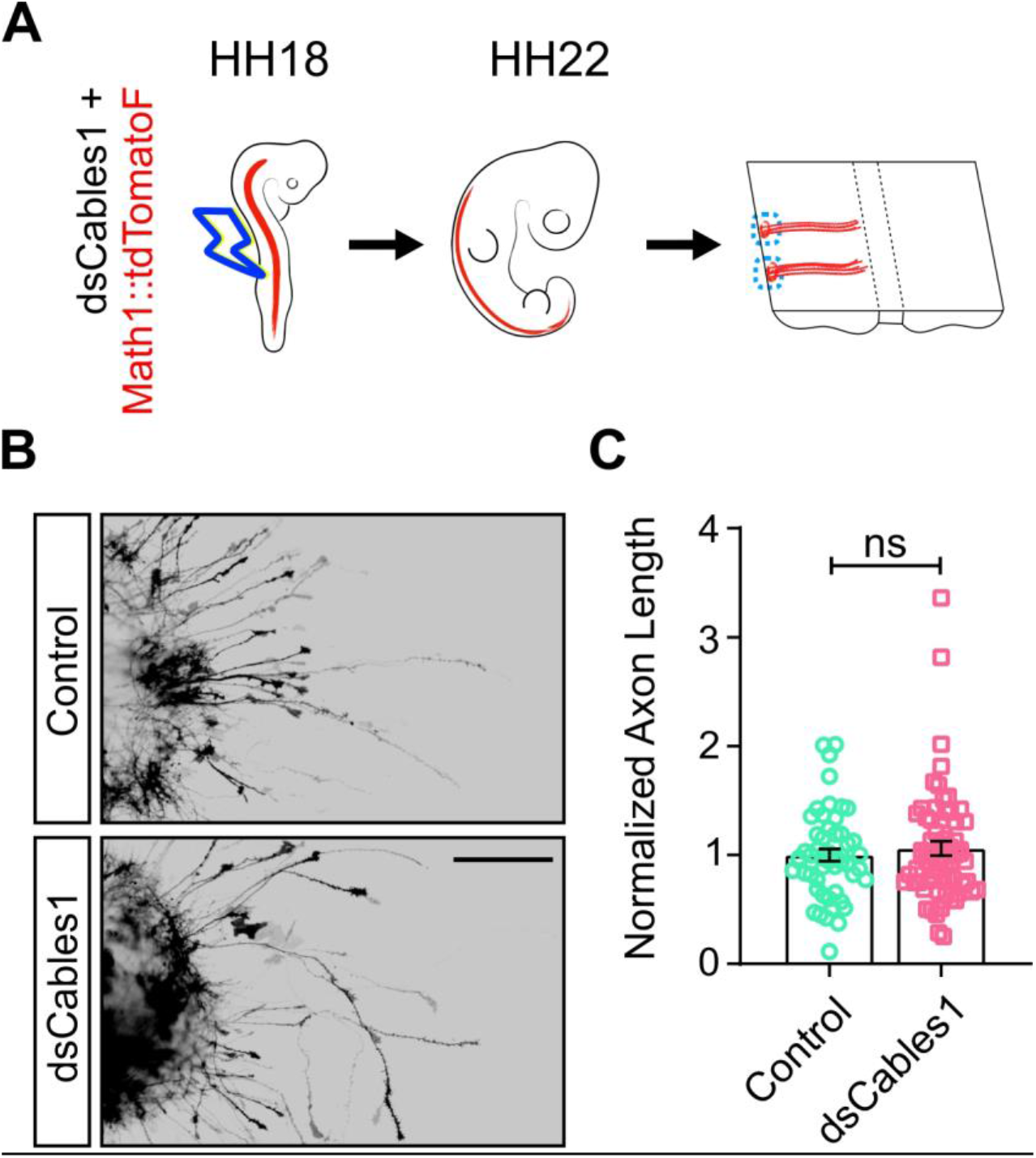
Cables1 is not required in pre-crossing axons. We excluded an effect of Cables1 on pre-crossing axons in vitro. HH18 embryos were injected and electroporated with Math1::td-Tomato-F (farnesylated td-Tomato under the control of the Math1 enhancer for specific expression in dI1 neurons) alone or together with dsRNA derived from Cables1 (dsCables1) (A). Explants of dorsal commissural neurons were prepared from open-book preparations of spinal cords dissected from HH21/22 embryos and cultured for 40 h. Axon outgrowth was visualized by staining for RFP (B). The average lengths of commissural axons were normalized to the lengths of control explants for each experiment. A total of 52 control explants (from 13 different embryos) and 64 explants taken from embryos electroporated with dsCables1 (n=16 embryos) from three independent experiments were quantified. The average length of neurites from control explants was 173 µm. See Methods for details. We did not find any significant difference (ns; p=0.4968) in neurite length (Unpaired T-test). Scale bar: 200 µm.

In contrast, when explants were taken from spinal cords dissected at HH26 (Figure 4A), axon lengths were significantly shorter from explants lacking Cables1 compared to control explants (Figure 4B,C). Taken together, our results suggested that Cables1 had an effect only on axons that had reached the floor plate, but not on pre-crossing axons. Therefore, we next tested whether Cables1 was involved in the responsiveness to Slit and Wnt. To this end, we added Slit2 or Wnt5a to the medium of commissural neuron explants dissected at HH26. As expected, axons from control explants were shorter in the presence of Slit2 (Figure 4B,C). However, no difference in axonal length was observed for neurons lacking Cables1 (Figure 4B,C). Adding Slit2 to the medium of neurons in the absence of Cables1 did not result in additional reduction of neurite length, indicating that Cables1 is required for the response to Slit2.

**Figure 4:**
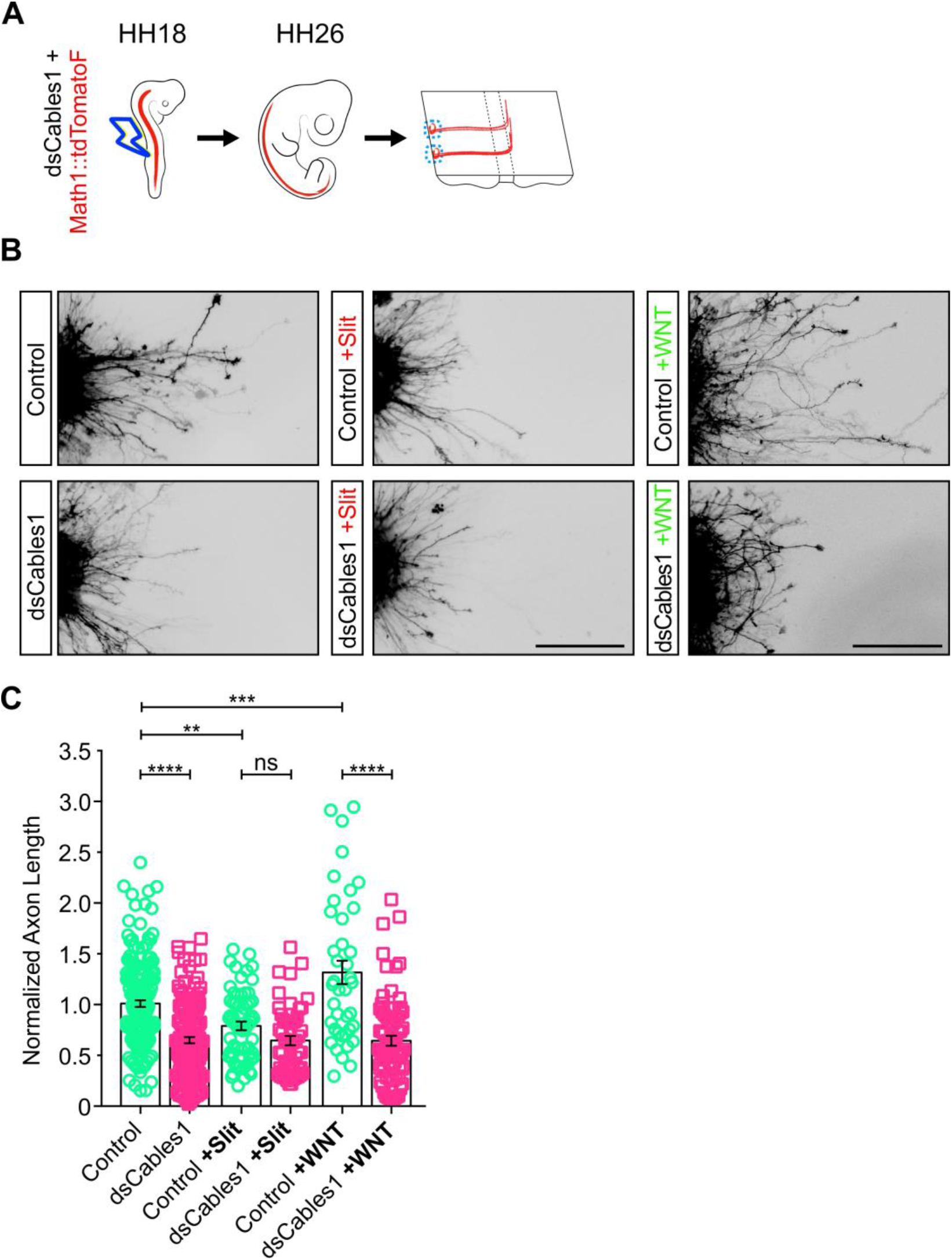
Loss of Cables1 prevents responsiveness to Slit and Wnt5a. Schematic representation of the experimental set-up (A). HH18 embryos were electroporated with Math1::td-Tomato alone or in combination with dsRNA derived from Cables1 (dsCables1). Explants of dI1 commissural neurons were prepared from open-book preparations of spinal cords dissected from HH26 embryos and cultured for 24 h before either medium (control), Slit2 (200 ng/ml), or Wnt5a (200 ng/ml) were added to the cultures for another 20 h. Explants were fixed and stained to visualize RFP (B). For each experiment, neurite lengths were measured and normalized to the length of control explants (C). Data from three different, independent experiments are pooled. Silencing Cables1 markedly reduced neurite lengths (n=37 explants) compared to controls (n=43). Adding Slit2 reduced neurite length of control explants (n=16), but not of explants lacking Cables1 (n=13). The lengths of neurites extending from control explants were increased after adding Wnt5a to the medium (n=10), but no increase in length was found after stimulating explants from embryos electroporated with dsCables1 with Wnt5a (n=20). ns, not significant; **p=0.0065; ***p=0.0007; ****p<0.0001; One-way ANOVA with Tukey’s multiple comparisons test. Scale bar: 200 µm.

Our previous studies demonstrated a growth-promoting effect of Wnt5a on post-crossing commissural axons (Avilés and Stoeckli, 2016). To demonstrate that this effect was also dependent on Cables1, we added Wnt5a to explants of commissural neurons dissected at HH26. In the presence of Wnt5a, axonal lengths of control but not explants lacking Cables1 was increased (Figure 4B,C). Taken together, these in vitro data confirm a requirement for Cables1 for both responsiveness to Slit and Wnt5a.

### Cables1 links Slit/Robo1 and Wnt/Fzd signaling *in vivo*

Our *in vitro* results were in agreement with the hypothesis that Cables was required for the responsiveness to Slit and the expulsion of axons from the floor plate, but also for Wnt-dependent turning of post-crossing commissural axons along the contralateral floor-plate border. To test this idea *in vivo*, we analyzed the functional interaction of Cables1 with Robo1 and β-Catenin, respectively. To this end, we used combinations of low doses of dsRNA targeting *Robo1, Cables1*, and *β-Catenin* that were not sufficient to induce aberrant phenotypes on their own. We reasoned that if these components interacted together in the same pathway, then combinatorial partial knockdown of these genes would result in aberrant axon navigation. We thus lowered the concentration of the dsRNA used for electroporation that effectively interfered with axon guidance (Supplementary Figure 4) to levels that were no longer inducing any changes in axonal behavior on their own (Figure 5). However, when we combined low concentrations of dsRNA targeting *Cables1* and *Robo1*, or *Cables1* and *β-Catenin*, we found significant effects on axon guidance (Figure 5B). The same was true for the combination of all three dsRNAs. Thus, these *in vivo* results confirmed a link between Slit/Robo and Wnt/Fzd signaling mediated by Cables1.

**Figure 5:**
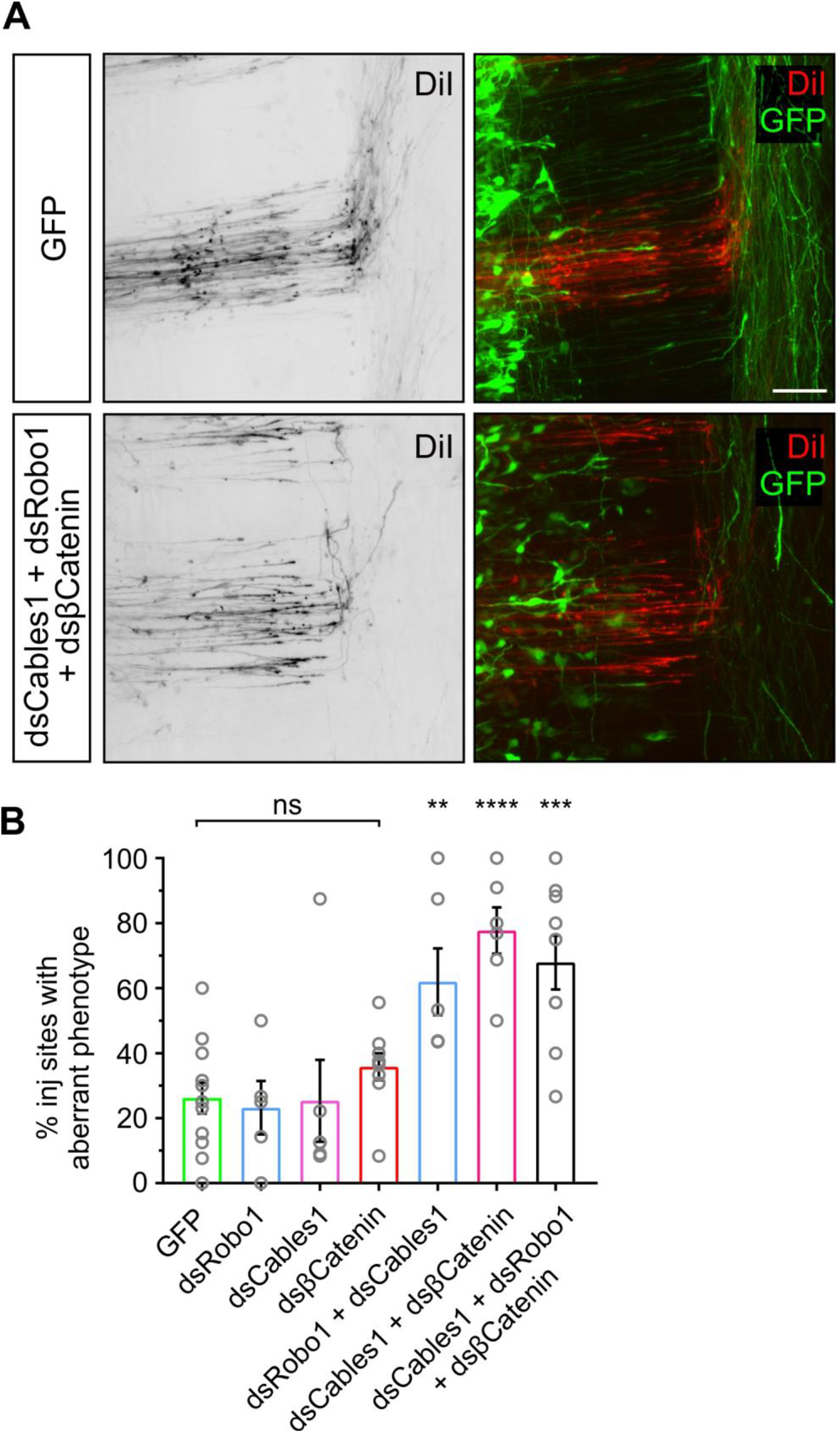
Cables1 cooperates with Slit/Robo and Wnt/Fzd signaling in axon guidance at the midline. Open-book preparations of control-treated (GFP-expressing) embryos and embryos electroporated with low concentrations of dsRNA derived from Cables1, Robo1, or β-Catenin alone or in combination were analyzed for axon guidance at the floor plate. In control-treated embryos, axons were crossing the midline and turning rostral upon floor-plate exit (A). In contrast, axons failed to turn correctly or did not reach the floor-plate exit site in embryos treated with combinations of any two low concentrations of dsCables1, dsRobo1, or dsβ-Catenin (not shown), or all three. (B) Compared to GFP-expressing control embryos, where an average of 26.2±4.8% of DiI injection sites with aberrant axonal trajectories were found (n=125, N=12), injection of only 75 ng/µl of dsRNA did not induce any aberrant axon guidance: dsRobo (23.2±8.2%; n=63 injection sites from N=5 embryos), dsCables1 (25.4±12.5%; n=56, N=6), dsβCatenin (35.8±4.1%; n=85, N=9). However, aberrant axonal navigation was found when combinations of dsRNA were used: dsCables1 + dsRobo1 (62.0±10.2%; n=85, N=6); dsCables1 + dsβCatenin (77.8±7.7%; n=70, N=6); dsCables1 + dsRobo1 + dsβCatenin (67.9±8.2%; n=92, N=9). ns not significant, **p<0.01, ***p<0.001, ****p=0.0001. One-way ANOVA with Dunnett’s multiple comparisons test. Scale bar: 50 µm.

### β-Catenin is preferentially phosphorylated in the post-crossing segment of commissural axons

To confirm the role of Cables1 as a linker between Slit/Robo and Wnt/Fzd signaling and to get detailed mechanistic insight, we analyzed the distribution of phosphorylated β-Catenin between pre- and post-crossing segments of commissural axons (Supplementary Figure 5). Abl kinase phosphorylates β-Catenin at tyrosine residue 489 (Rhee et al., 2007). Staining with a pY489-specific antibody revealed an accumulation of β-Catenin pY489 in the distal, post-crossing axonal segment both *in vivo* (Supplementary Figure 5C,D) and *in vitro* (Supplementary Figure 5F).

Based on our results demonstrating that Cables1 had an effect on post-but not pre-crossing axons, we compared the localization of total β-Catenin and β-Catenin pY489 in neurons dissected from HH21 and HH26 embryos (Figure 6). We found no difference in levels of total β-Catenin between proximal and distal segments of pre-crossing or post-crossing dI1 axons (Figure 6B,C). However, β-Catenin pY489 levels were higher in distal segments of post-crossing axons (Figure 6D,E). No such difference between proximal and distal axonal segments was seen for pre-crossing axons.

**Figure 6.**
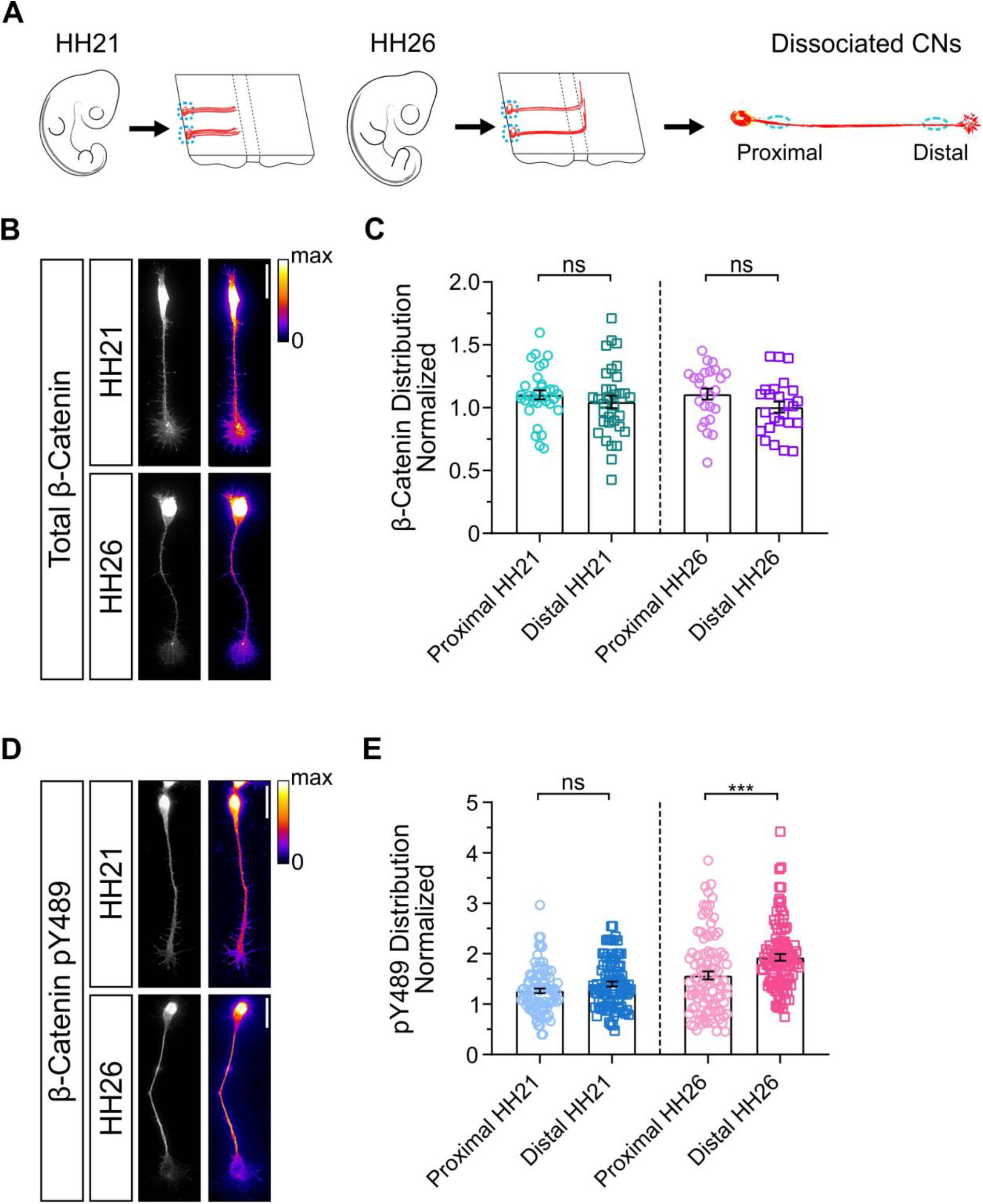
β-Catenin pY489 accumulates in the distal part of post-crossing commissural axons. (A) Schematic representation of the experimental design. Dissociated commissural neurons were prepared from open-book preparations of HH21 and HH26 embryos, grown for 40-48 h before staining with antibodies recognizing β-Catenin or specifically β-Catenin pY489. Fluorescence intensity in the proximal and distal parts of the axons were measured and normalized to the average intensity of the entire axon. Total levels of β-Catenin measured at HH21 and HH26 did not differ between the proximal and the distal axon (B,C; HH21, n=33 neurons; HH26, n=25 neurons). Similarly, comparing levels of β-Catenin pY489 between proximal and distal axons of axons collected from HH21 embryos did not differ (D,E). In contrast, levels of β-Catenin pY489 were significantly higher in distal compared to proximal axons of neurons collected from HH26 embryos (D,E; HH21, n=95 neurons; HH26, n= 99 neurons; p=0.0538 for HH21 and ***p=0.0009 for HH26). Results were obtained from three independent experiments. ns=not significant; Paired t-test.

Next, we looked at the distribution of β-Catenin pY489 in neurons lacking Cables1 (Figure 7). The observed accumulation of β-Catenin pY489 in the distal axon disappeared in the absence of Cables1, indicating that Cables1 was responsible for the accumulation of β-Catenin pY489 in the distal post-crossing axons.

**Figure 7.**
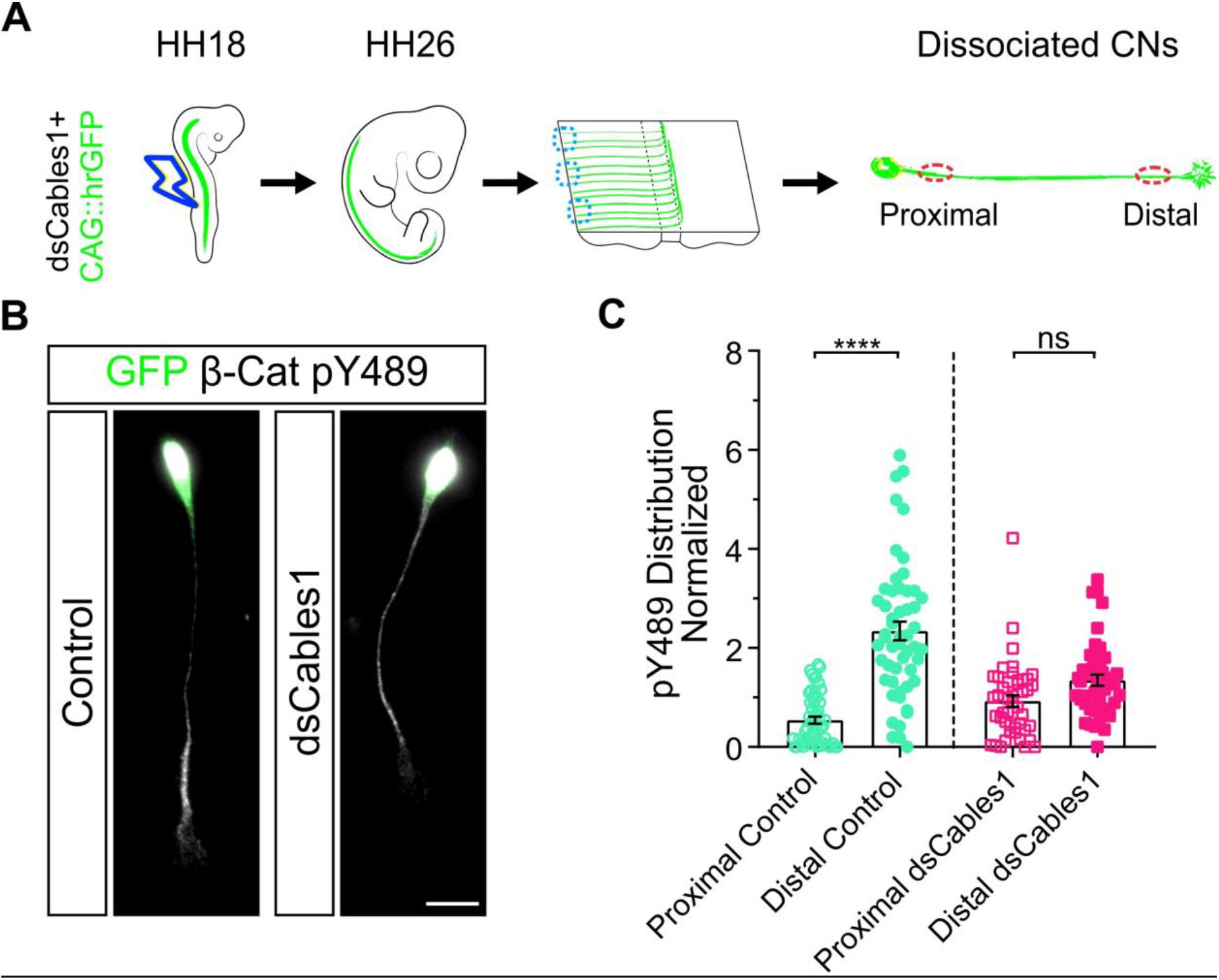
Cables1 regulates the phosphorylation of β-Catenin Y489 in post-crossing commissural axons. (A) Schematic of the experimental design. HH18 embryos were electroporated with dsRNA derived from Cables1 and a GFP reporter plasmid. Dissociated commissural neurons were prepared from open-book preparations of HH26 spinal cords, stained with an antibody specific for β-Catenin pY489 and used for intensity measurements. (B) Immunostaining of commissural neurons for β-Catenin pY489 (white) and GFP (green) isolated from control-treated and dsCables1-electroporated embryos demonstrated that the accumulation of β-Catenin pY489 depended on Cables1, as intensity levels did no longer significantly differ when we compared distal and proximal axons from embryos lacking Cables1. (C) Quantification of β-Catenin pY489 levels in proximal compared to distal axons in control and dsCables1-transfected neurons normalized to the average intensity of the entire axons (±s.e.m.). Results were obtained from three independent experiments. Control, n=53 neurons; dsCables1, n=45 neurons. Control, ****p<0.0001; dsCables1, ns, p=0.1382; Tukey’s multiple comparisons test.

### Phosphorylation of β-Catenin at position Y489 is required for post-crossing axon growth

Next, we wanted to test our model that Cables1-mediated localization of phosphorylated β-Catenin in the distal axon/growth cone was required for axon guidance *in vivo*. To this end, we generated two different β-Catenin mutants: β-CateninY489E, a phosphomimetic (constantly active) mutant, and β-CateninY489F, a mutant that cannot be phosphorylated (non-active form) (Figure 8). Both mutant forms of β-Catenin were specifically expressed in dI1 neurons with the help of the Math1 enhancer. Aberrant axonal turning into the longitudinal axis in the absence of Cables1 could be rescued by co-expression of β-CateninY489E, the constantly active form of β-Catenin, but not with the non-active mutant of β-Catenin, β-CateninY489F (Figure 8). The overexpression of each of these mutant versions of β-Catenin, Math1::β-CateninY489E and Math1::β-CateninY489F in the presence of Cables1 did not result in axon guidance phenotypes per se (Supplementary Figure 6).

**Figure 8.**
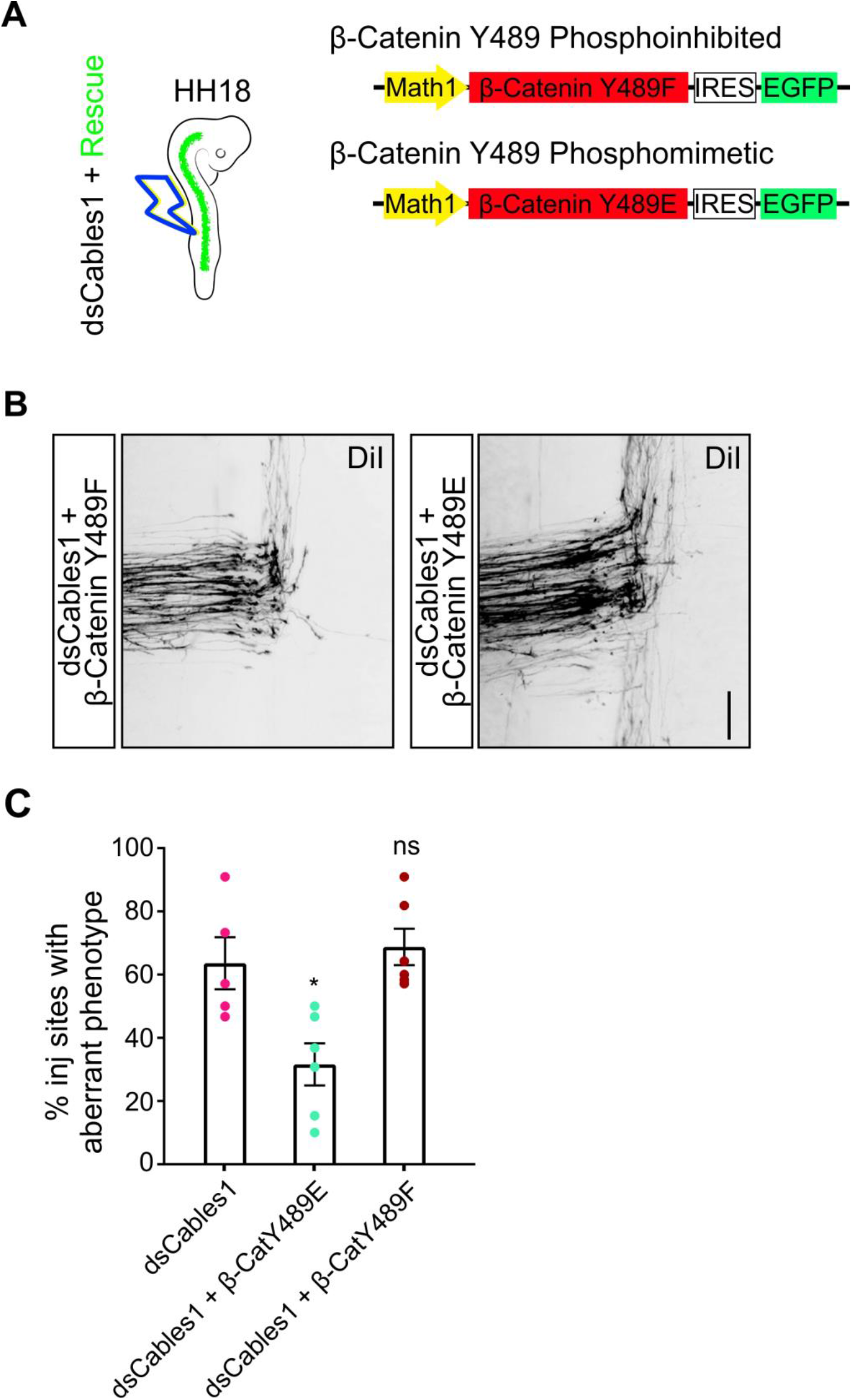
A phosphomimetic version of β-Catenin pY489 can rescue loss of Cables1 function. (A) Schematic representation of the constructs used in this study. Embryos were electroporated at HH18 with dsRNA derived from Cables1 in combination with mutant versions of β-Catenin. Axon guidance phenotypes induced by the absence of Cables1 function could be rescued with the dominant active (phosphomimetic) version of β-Catenin (β-Catenin Y489E) but not with β-Catenin Y489F, which cannot be phosphorylated (B). (C) Quantification of aberrant axon guidance phenotypes obtained from rescue experiments. Only co-electroporation of Math1::β-Catenin Y489E together with dsCables1 was able to reduce the percentage of DiI injection sites with aberrant axon guidance phenotypes obtained upon loss of Cables1 function. We saw an average of 63.6±8.2% of the DiI injections sites with aberrant axon guidance in embryos electroporated with dsCables1 (n=65 injection sites in N=5 embryos). The percentage of DiI injection sites with aberrant axonal trajectories was reduced to 31.6±6.6% by co-electroporation with β-Catenin Y489E (n=80, N=6). No rescue was obtained after co-electroporation with the β-Catenin mutant that could not be phosphorylated, β-Catenin Y489F (68.7±5.7% injection sites with aberrant axon guidance, n=82, N=6) *p=0.0147; ns= not significant. Dunnett’s multiple comparisons test. Scale bars: 50 µm.

Taken together, our *in vivo* and *in vitro* data are in line with the model that Cables1 links Slit/Robo signaling during midline crossing with Wnt/Fzd signaling required for post-crossing axons to turn into the longitudinal axis of the spinal cord. Cables1 is required for the localization of phosphorylated β-Catenin pY489 in distal axons and growth cones, which in turn is required for correct turning of post-crossing commissural axons along the antero-posterior Wnt gradient.

## DISCUSSION

For axonal navigation towards the intermediate target, arrival and departure without lingering, and continuation towards the next intermediate or the final target, the changes in receptor expression on the growth cone surface have to be orchestrated in a precisely controlled manner. Most studies have concentrated on the regulation of receptor expression of a particular signaling pathway. In contrast, this study describes how two known signaling pathways are connected. Using *in vivo* and *in vitro* assays, we identified a role for Cables1 as a linker between the Slit/Robo-mediated floor-plate exit of commissural axons and their Wnt/Fzd-mediated turn into the longitudinal axis. Previous studies demonstrated that expression of Robo1, the receptor for the repulsive Slit molecules expressed by the floor plate, is regulated at the post-translational level (Alther et al., 2016; Pignata et al., 2019; Kinoshita-Kawada et al., 2019). This mechanism is in line with descriptions in flies, where Robo expression was shown to depend on trafficking as well (summarized by Gorla and Bashaw, 2020). In vertebrates, miRNA-mediated regulation of translation has been identified as additional regulatory mechanism of Robo surface expression (Yang et al., 2018).

Responsiveness to the guidance cues regulating axon guidance along the longitudinal axis of the spinal cord is controlled by different mechanisms. The expression of Hhip (Hedgehog-interacting protein), the Shh receptor on post-crossing axons, is regulated at the transcriptional level by Shh itself in a Glypican-1-dependent manner (Bourikas et al., 2005; Wilson et al., 2013). In contrast, expression of Fzd3, the Wnt receptor on post-crossing axons, is also regulated by specific trafficking (Alther et al., 2016; Onishi and Zou, 2017). Robo and Fzd receptors are transported to the growth cone surface in a Calsyntenin1-dependent manner but in different vesicles (Alther et al., 2016). Timing of vesicular transport of Robo1 and its insertion into the growth cone membrane has been shown to depend on RabGDI (Philipp et al., 2012; Alther et al., 2016). The timer of Fzd3 transport and insertion is still elusive, as RabGDI is not involved in Fzd3 expression.

In addition to the controlled delivery of guidance receptors, a molecular linker between the Slit/Robo and the Wnt/Fzd pathway provides a regulatory mechanism for fine-tuning the temporal sequence of events and to link floor-plate exit with turning of post-crossing axons. Our results demonstrate that Cables1 is required for axon guidance (Figure 2) rather than just axonal growth (Figure 3 and Supplementary Figure 3) or neuronal differentiation (Supplementary Figure 2), although growth and guidance cannot be separated for post-crossing axons (Figures 2 and 4). Post-crossing axons are shorter *in vitro* and fail to respond to Wnt5a in the absence of Cables1 (Figure 4).

Our results are in line with a model that suggests an association between Abl and Robo1 in axons crossing the midline (Rhee et al., 2007, but see also Bashaw et al., 2000). Upon Slit binding, Robo1 receptors are internalized and the associated Abl molecule is detached from Robo via Cables1. Cables1 brings Abl in close proximity to β-Catenin, which gets phosphorylated at tyrosine residue 489. This phosphorylation changes the interactions of β-Catenin and prepares it for its role in Wnt signaling at the floor-plate exit site.

Our study is the first report on the involvement of a Robo/Cables1/β-Catenin link in commissural axon guidance. So far, Robo-mediated expulsion of axons from the Slit expressing floor plate has not been functionally linked to guidance cues for the longitudinal axis. We previously demonstrated a role of β-Catenin in Wnt signaling and guidance of post-crossing commissural axons (Avilés and Stoeckli, 2016). Here, we extend these findings and demonstrate that β-Catenin needs to be phosphorylated at tyrosine 489 for its role in post-crossing axon guidance (Figure 8). This finding is intriguing in the context of a recent study about axonal navigation at the chiasm (Morenilla-Palao et al., 2020). The authors found phosphorylation of β-Catenin at tyrosine 654 in ipsilaterally projecting retinal ganglion cell axons. In contrast to Abl-mediated β-Catenin phosphorylation in post-crossing dI1 axons, which is on tyrosine 489, ipsilaterally projecting axons in the visual system are phosphorylated by EphB1. In both cases, crossing the chiasm and crossing the floor plate requires β-Catenin, as silencing β-Catenin resulted in axonal stalling at the midline (Morenilla-Palao et al., 2020; Avilés and Stoeckli, 2016, this study).

Taken together, our *in vivo* and *in vitro* results suggest a model for Cables1 function that connects the Robo1-mediated exit from the floor plate in response to Slit binding to the attractive effect of Wnts directing post-crossing axons rostrally. Cables1 links Robo1-bound Abl kinase to β-Catenin. The accumulation of Abl-dependent phosphorylation of β-Catenin at tyrosine 489 in the growth cone/distal axon is required for post-crossing axons to respond to Wnt5a. Our results demonstrate that Cables is required for the distal localization of phosphorylated β-Catenin-pY489 (Figure 7). In turn, co-electroporation of a dominant active form of β-Catenin-pY489, β-Catenin-Y489E, can rescue the lack of Cables1 (Figure 8), indicating that β-Catenin-pY489 is required for Wnt responsiveness of post-crossing axons upon floor-plate exit.

## EXPERIMENTAL PROCEDURES

### Animals

Fertilized chicken eggs were obtained from a local supplier and incubated at 39°C. All the experiments including chicken embryos were carried out in accordance with Swiss law on animal experimentation and approved by the cantonal veterinary office of Zurich.

### Method details

#### In ovo electroporation

After 2 or 3 days of incubation at 39°C, fertilized eggs were windowed for injection and electroporation, as described previously (Wilson and Stoeckli, 2011, and 2012). Embryos were staged according to Hamburger and Hamilton (1992). Unilateral electroporations were performed at embryonic day 3, HH17-18, using 5 pulses of 25 Volts of 50 msec duration and 1 sec interpulse interval. For bilateral electroporation or electroporation at E2, we used 5 pulses of 18 Volts only.

#### Plasmids and dsRNA

For functional gene analysis, chicken embryos were injected and electroporated with long dsRNA (300 ng/µl) derived from the target gene and a plasmid encoding β-actin-driven hrGFP (25 ng/µl). For hypomorphic experiments, combination of low doses (75 ng/µl) of each dsRNA were used.

For the pY489 phosphomutant versions of β-Catenin, we used the Q5 Site-directed mutagenesis kit (NEB) to generate β-CateninY489F, a form of β-Catenin that cannot be phosphorylated at tyrosine 489 due to the exchange of tyrosine 489 with phenylalanine, and β-CateninY489E (exchange of tyrosine 489 with glutamic acid), a phosphomimetic form of β-Catenin that is dominant active. The following primers were used for the phosphomimetic substitution of Tyr (tat) by Glu (caa): Fw: 5’-TCGCCTTCATcaaGGACTGGCCTGTTG-3’ Rv: 5’-ACGGCATTCTGGGCCATC-3’. For the phosphoinhibited substitution of Tyr (tat) by Phe (ttt): Fw: 5’-TCGCCTTCATtttGGACTGCCTG-3’ Rv: 5’-ACGGCATTCTGGGCCATC-3’. For rescue experiments, the ORF of mouse Cables1 was obtained from Biocat/Origene. The amplified PCR fragment was subcloned via HIFI cloning (NEB) under a Math1 enhancer.

#### Open-book preparations and DiI tracing

Spinal cords were dissected at HH25-26 (E5) as open-book preparations and fixed for 30 min in 4% PFA in PBS. To label dI1 commissural neurons, we injected Fast-DiI (5 mg/ml in ethanol; Thermo Fisher) into the area of the cell bodies in the dorsal spinal cord, as described previously (Wilson and Stoeckli, 2012; Perrin and Stoeckli, 2000). The phenotypes were analyzed by an observer blind to the experimental condition and categorized as follows: normal (axons cross the floor plate and turn rostrally into the longitudinal axis along the contralateral floor-plate border), floor-plate stalling (more than 50% of the axons fail to cross the floor plate), or no turning (more than 50% of the axons that reach the contralateral floor-plate border fail to turn into the longitudinal axis). For embryos dissected at HH29-30 (E6), ipsilateral turns were excluded from the quantification, as later developing populations of axons labelled by late DiI injections normally extend to the floor plate without crossing. Therefore, these normally navigating axons could not be distinguished from dI1 axons stalling at the floor-plate entry site. Images were acquired using an Olympus BX61 microscope equipped with a spinning disk unit. Data are given as mean±sem (standard error of the mean). Statistical analysis was performed using Prism 8 (GraphPad).

#### In situ hybridization and immunostaining

Embryos were sacrificed and fixed in 4% paraformaldehyde in PBS at room temperature for different times depending on the stage. The tissue was cryoprotected by incubation in 25% sucrose/PBS and then embedded in Tissue-Tek O.C.T Compound (Sakura). Specimens were frozen in isopentane on dry ice and stored at −20°C. Sections of 25 μm thickness were obtained using a cryostat (LEICA CM1850). Expressed sequence tags (ChESTs; SourceBioScience) were used to generate *in situ* probes using a DIG RNA labeling kit (Roche). In situ hybridization was performed as described (Mauti et al., 2006). Immunostaining was performed as described previously (Perrin et al., 2001; Wilson and Stoeckli, 2011). The complete list of ChESTs and antibodies can be found in Table 1 (Key Ressources Table). For immunostaining, cultures of dissociated neurons and explants were fixed in 4% PFA for 15 minutes at room temperature. Cells and explants were incubated in 100 mM glycine for 20 minutes and permeabilized with PBST (0.25% Triton X-100 in PBS) for 15 min. To reduce unspecific binding of antibodies, cells/explants were incubated in 10% FCS in PBST at room temperature for 30 min. Primary antibodies diluted in 10% FCS/PBST were incubated at 4°C overnight. The following day, cultures were washed twice in PBST and incubated with secondary antibodies diluted in 10% FCS/PBST for 2 hours at room temperature. Before mounting in Mowiol-DABCO, samples were washed three times in PBS.

#### Quantitative Real-Time-PCR

RNA was isolated from spinal cords of embryos at HH22 and HH25 using the RNeasy mini kit (#74134, QIAGEN). For the evaluation of RNAi efficiency, chicken embryos were electroporated at HH17-18 and neural tubes were dissected 24 hours later under a fluorescence microscope (Olympus SZX12). Total RNA was then reverse transcribed using SuperScript(tm) III First-Strand Synthesis SuperMix (#18080-400, Thermo Fisher). The primers used for the quantitative Real-Time PCR reaction are listed in Table 1 (Key Resources Table). qRT-PCR was performed using the Fast Sybr Green Master Mix (#4385610, Thermo Fisher) and run on a QuantStudio 3 Real Time PCR System (Applied Biosystems). mRNA expression levels were normalized to the expression level of chicken 18S ribosome (Himmels et al., 2017) and quantified using the 2^-ΔΔCt^ method. PCR amplifications were assessed from pools of spinal cords from at least three independent experiments. For quantification of *Cables1* isoform levels and *Cables2*, values were normalized to HH22 *Cables1_X1*.

#### SDS-PAGE and Western Blotting

For the evaluation of RNAi efficiency, chicken embryos were electroporated at HH17-18 with a combination of two different dsRNAs targeting Cables1 (300 ng/µl each). Neural tubes were dissected 48 hours later under a fluorescence microscope (Olympus SZX12). Cells were lysed with RIPA buffer (150 mM NaCl, 1% Nonidet P-40, 0.5% Sodium deoxycholate, 0.1% SDS, 50 mM Tris-HCl, pH 7.4) supplemented with protease inhibitors (Roche 11836170001) and phosphatase inhibitors (5 mM NaF, 1 mM Na_3_VO_4_, 10 mM β-Glycerophosphate). Protein concentrations were measured and samples were prepared for PAGE by adding 0.2 volumes of 5X Loading Buffer (650 mM Tris-Cl, pH 6.8, 5 % SDS, 25 % Glycerol, 500 mM DTT and bromophenol blue) and incubated for 5 min at 95°C. Protein samples were separated by SDS-PAGE and transferred to a PVDF membrane. The membranes were blocked with 5% milk in TBST (0.01 M Tris-HCl, pH 7.5, 150 mM NaCl, 0.1% Tween20), followed by primary antibody incubation overnight at 4°C. On the following day, membranes were washed in TBST three times for 15 min, before incubation for 2 h at room temperature with the corresponding secondary antibodies conjugated to horseradish peroxidase. Membranes were washed in TBST before using the ECL Western Blotting Detection Reagent (GE Healthcare). The chemiluminescence signal was detected using the Amersham Imager 600 (GE Healthcare).

#### Quantification of axonal outgrowth from commissural explants

For average length measurements, only td-Tomato-F-positive dI1 axons (labelled by electroporation of embryos with Math1::td-TomatoF) were used. Each explant was divided into 4 quadrants and the average neurite length from the explant border was measured for each quadrant with ImageJ (v 1.52i, Java 1.8.0_101 64-bit, NIH). Axons were manually traced using a Wacom DTU-1931 tablet and pen tool. Data from at least three different, independent experiments were pooled and normalized to control conditions of each independent experiment. Data are given as mean±sem. Statistical analysis was performed using Prism 8 (GraphPad).

For neurite measurements in half open-book explants, axon outgrowth was quantified with a Scholl analysis using the ImageJ plug-in Neurite-J (Torres-Espín et al., 2014). Data were plotted as the number of neurites (y-axis) crossing a concentric ring around the explant with the given distance (x-axis) from explant border.

#### Quantitative Analysis of phospho-Y489 levels in commissural neurons

To analyze total β-Catenin and β-Catenin-pY489 levels in commissural neurons, we performed fluorescence intensity measurements with ImageJ (v 1.52i, Java 1.8.0_101 64-bit, NIH). Axons were carefully delineated (excluding the growth cone) to acquire mean levels of fluorescence. For comparison of axonal distribution, we measured the mean fluorescence in a determined area in the distal and proximal axon and normalized to total axon levels. To account for background signal, we measured the mean fluorescence value by selecting an area adjacent to the axon. For quantification, at least 20 neurons per condition were measured using a Wacom DTU-1931 tablet and pen tool. Data from at least three different, independent experiments are pooled and normalized to control conditions of each independent experiment. Data are given as mean±sem. Statistical analysis was performed using Prism 8 (GraphPad).

#### Primary Neuron Cultures and Explant Cultures

Explants of commissural neurons were obtained from dorsal spinal cords dissected from HH25-26 embryos for post-crossing, and HH22 for pre-crossing neurons. To ensure that we dissected dI1 commissural neurons, embryos were dissected under a fluorescent stereoscope in order to visualize Math1::td-TomatoF-positive cells. Commissural explants were grown on 8-well LabTek slides (Nunc) coated with poly-Lysine (20 μg/ml; Sigma) and Laminin (10 μg/ml). The medium for commissural neurons was as previously described (Niederkofler et al., 2010), except for pre-crossing cultures, where the medium was supplemented with recombinant Netrin (100 ng/ml, R&D Systems).

For experiments assessing Slit and Wnt responsiveness, control medium or medium containing Slit2 (200 ng/ml; R&D Systems) or Wnt5a (200 ng/ml; R&D Systems) was added to the commissural neurons after 48 hours or to the explants after 24 hours in vitro. Explants were grown for additional 20 hours before fixation and immunostaining.

For half open-book explants, spinal cords were dissected from HH25-HH26 embryos as open-book preparations. The electroporated half of the spinal cord including the floor plate was dissected, cut in pieces and cultured on 8-well LabTek slides (Nunc) coated with poly-Lysine (20 μg/ml; Sigma) and Laminin (10 μg/ml). The culture medium was as previously described (Niederkofler et al., 2010). Explants were grown in vitro for 28 hours before fixation and immunostaining.

For cultures of dissociated commissural neurons, neurons were obtained from dorsal spinal cords dissected from HH25-26 embryos for post-crossing, and HH21 for pre-crossing cultures. Commissural neurons were grown on 8-well LabTek slides (Nunc) coated with poly-L-Lysine (20 μg/ml; Sigma) and Laminin (10 μg/ml). The culture medium was as previously described (Niederkofler et al., 2010), except for pre-crossing cultures where the medium was supplemented with recombinant Netrin (50 ng/ml, R&D Systems). Primary neurons were plated at low density (8’000-10’000 cells/well) and kept in an incubator with 5% CO_2_ at 37°C. Cultures were grown for 40-48 h before fixation and immunostaining.

## KEY RESOURCES TABLE

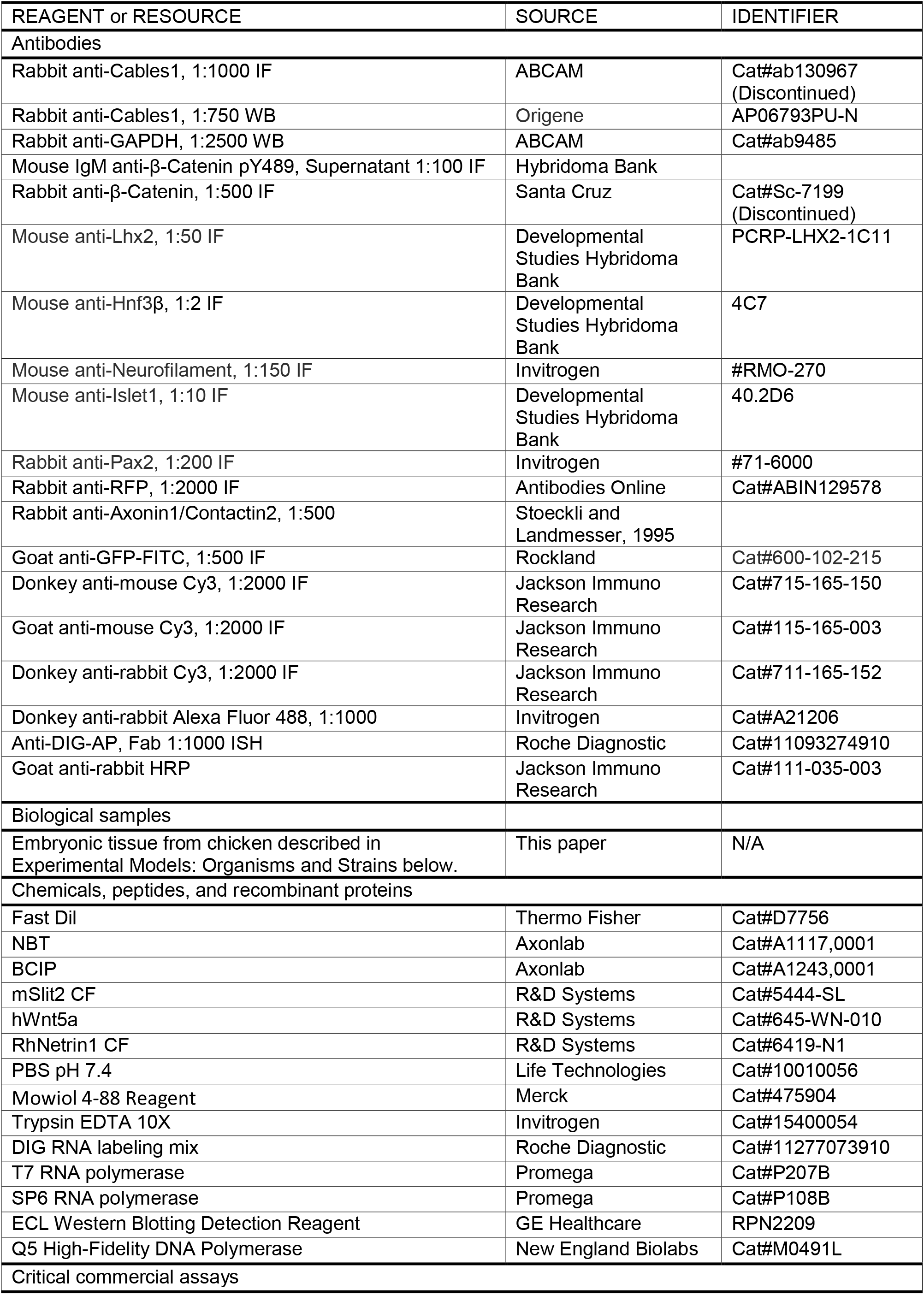
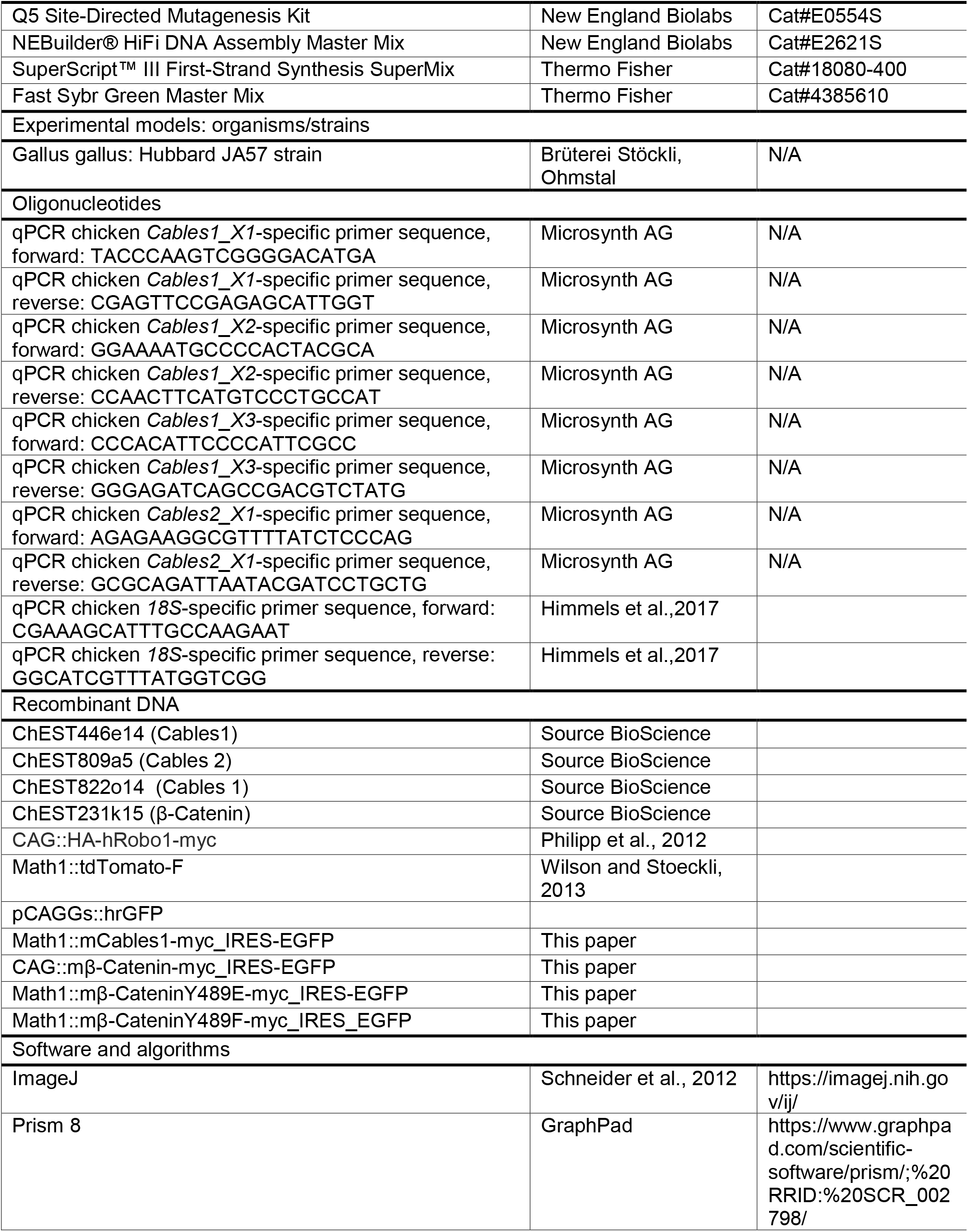
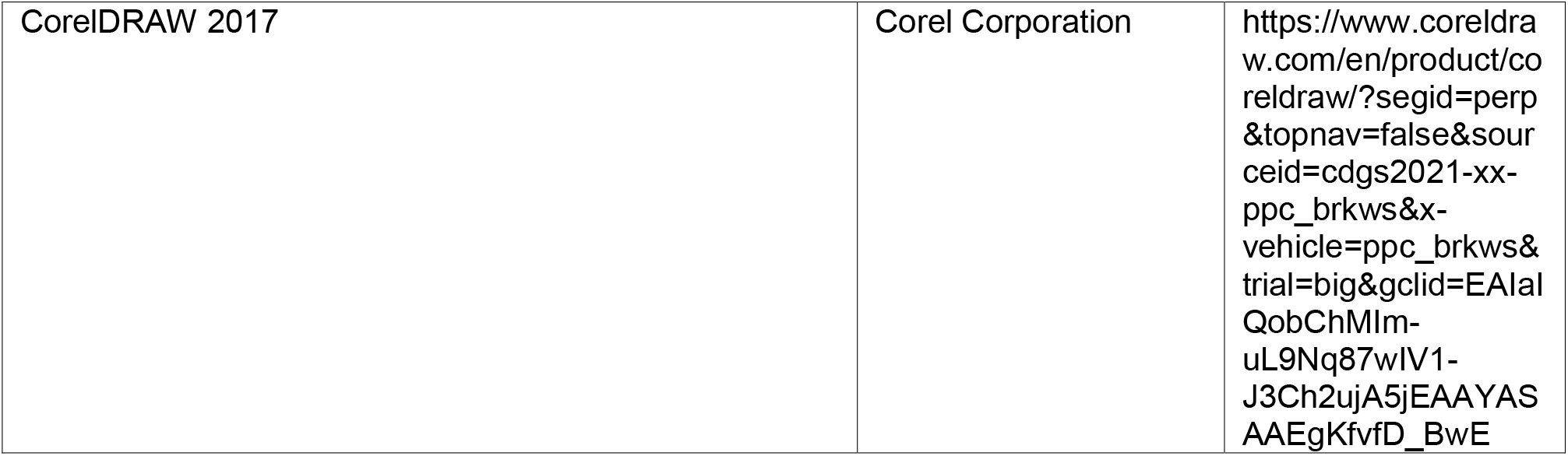

## ACKNOWLEDGEMENTS

We thank Tiziana Flego and Dr. Beat Kunz for excellent technical assistance. This project was supported by the Swiss National Science Foundation.

## Supplementary Figures

**Supplementary Figure 1:**
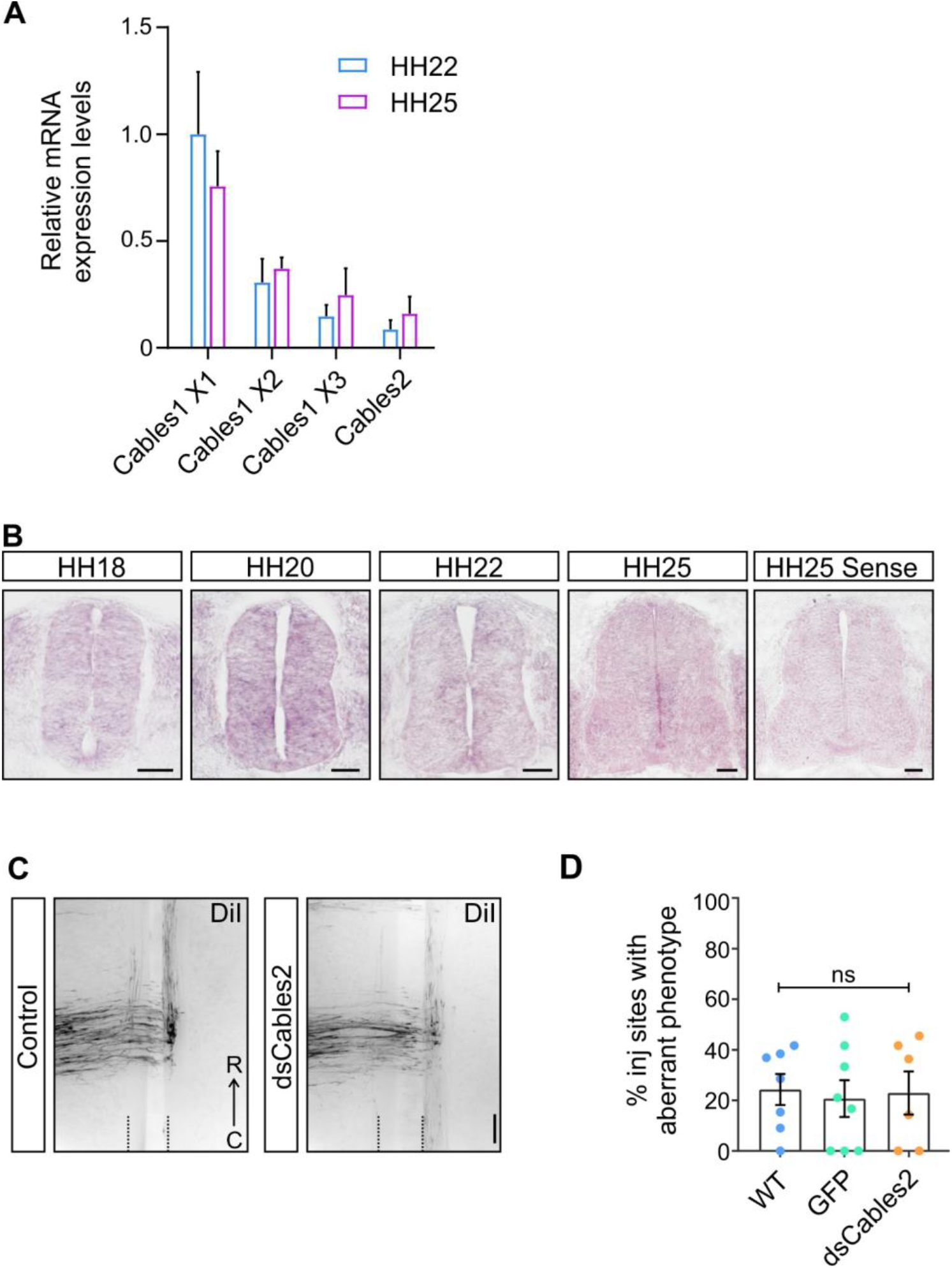
In the developing spinal cord Cables2 is expressed at very low levels, if at all, and is not involved in commissural axons guidance at the midline. qRT-PCR analysis showed high expression levels of *Cables1* isoformX1 in comparison to *Cables1* isoformX2 and isoformX3, as well as *Cables2* at stages HH22 and HH25. All mRNA levels were normalized to *Cables1* isoformX1 at HH22 (A). *Cables2* mRNA was found at low levels throughout the developing neural tube during the time window, when dI1 commissural axons cross the floor plate and turn rostral along the contralateral floor-plate border (compare sections hybridized with anti-sense probe to the section hybridized with the sense probe). In contrast to what we observed for *Cables1* mRNA (Figure 1), *Cables2* mRNA was not upregulated in dI1 commissural neurons during the time when their axons cross the midline (B). Functional analysis of Cables2 revealed no effect on dI1 axon guidance at the floor plate (C). Quantification of aberrant commissural axon trajectories in non-injected control embryos (WT, non-treated controls, 24.3±6.1%; n=92 injection sites in N=7 embryos, same group as shown in Figure 2), GFP-expressing control embryos (GFP; 20.7±7.2%; n=93, N=8), and embryos electroporated with dsRNA derived from Cables2 (dsCables2; 23.0±8.4%; n=55, N=6) showed no significant (ns) difference in the number of DiI injection sites with aberrant axonal trajectories (D). Values are given as mean ± s.e.m (D). Bar: 50 µm.

**Supplementary Figure 2.**
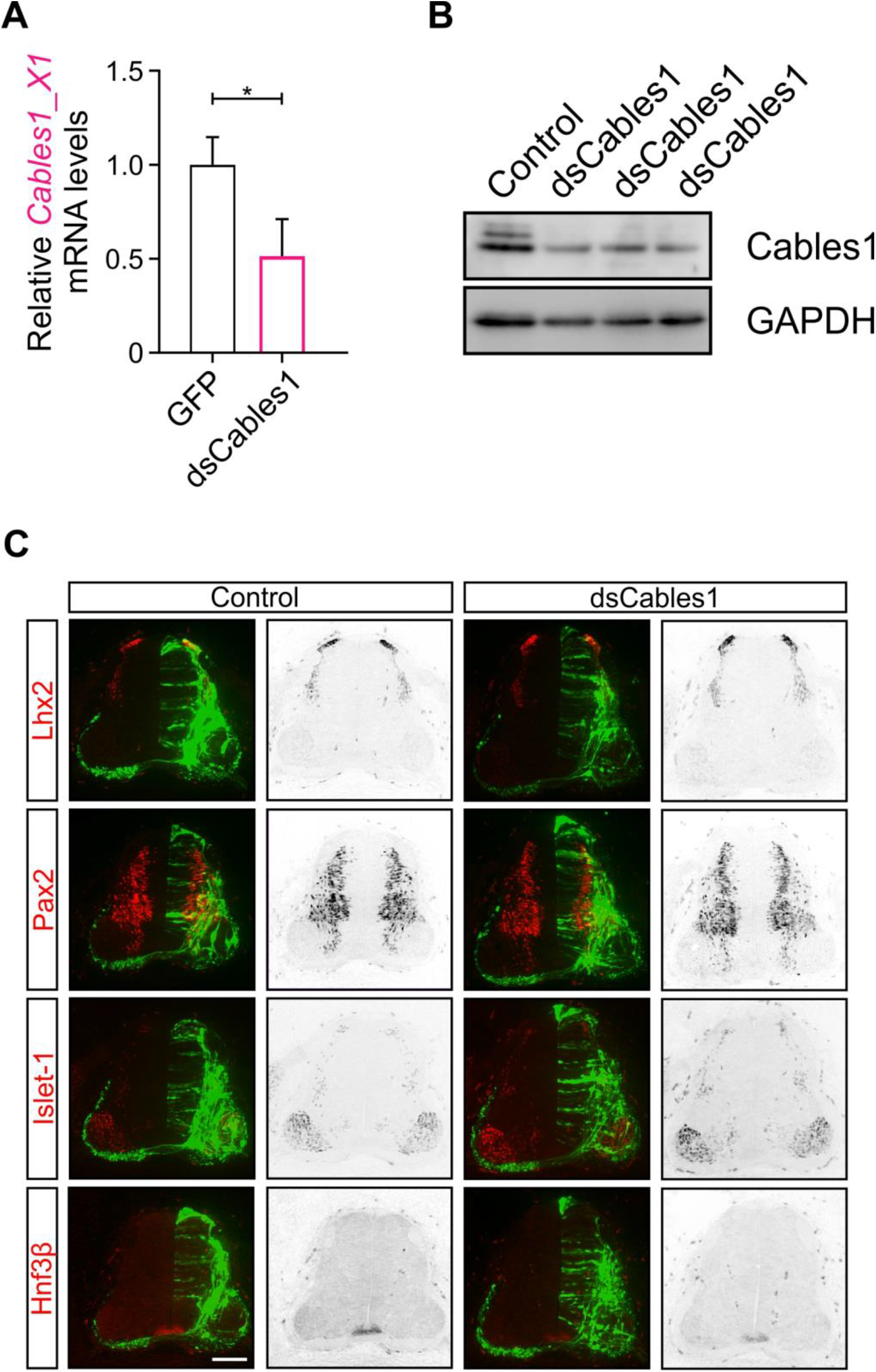
Downregulation of Cables1 by in ovo RNAi effectively reduces levels of Cables1 and does not interfere with neuronal differentiation. Effective downregulation of Cables mRNA was demonstrated with qRT-PCR for isoform X1 (A) with 4 pools of embryos. *p=0.03, paired t-test. Western Blotting of proteins isolated from HH25 spinal cords confirmed effective downregulation of Cables1 in three independent pools of embryos electroporated with dsCables1 at HH17/18 (B). Levels were reduced by 50-60%. With the parameters used to silence Cables1, we successfully transfected on average 50% of the cells in the targeted area of the neural tube. Therefore, the observed reduction in Cables1 protein indicate a more or less complete removal of Cables1 from the transfected cells. Immunostaining of HH25 spinal cord sections with antibodies against Lhx2 (marker for dI1 neurons), Pax2 (interneurons), Islet-1 (motoneurons), and Hnf3β (floor-plate cells) did not reveal any differences in neuronal differentiation between control-treated, GFP-expressing embryos, and experimental embryos electroporated with dsRNA derived from *Cables1* at HH18 (C).

**Supplementary Figure 3:**
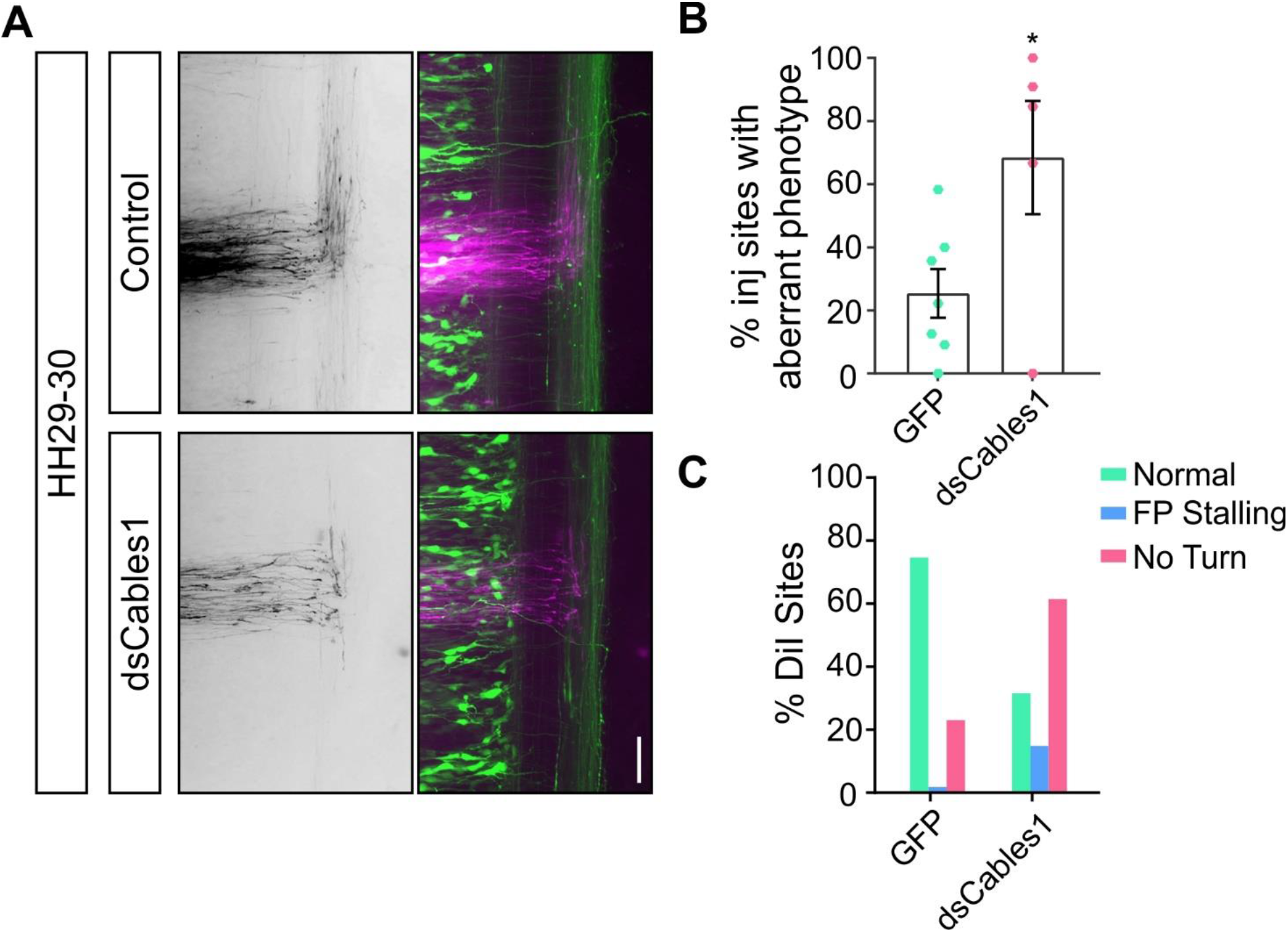
The observed axon guidance defects after silencing Cables1 are not explained by a reduction in neurite growth speed. To exclude a slower growth rate as the reason for the failure of axons to turn into the longitudinal axis in the absence of Cables1, we analyzed dI1 axonal navigation at the floor plate in open-book preparations of spinal cords dissected from embryos at HH29/30, that is 1.5 days older than those shown in Figure 2. The fact that axons still stalled at the floor-plate exit site and failed to turn into the longitudinal axis confirmed that axon guidance defects could not be explained by a slower growth rate, but had to be due to a failure to respond to guidance cues provided by the floor plate (A). When compared to control-treated embryos, aberrant axon guidance was found at 68.4±17.9% of the DiI injection sites in embryos electroporated with dsCables1 (n=56 injection sites in N=5 embryos), compared to control-injected embryos (GFP plasmid only), where aberrant axonal trajectories were seen only at 25.4±7.7% of the injection sites (* p=0.0338; n=74 injection sites in 7 control-injected embryos). Scale bar: 50 µm.

**Supplementary Figure 4:**
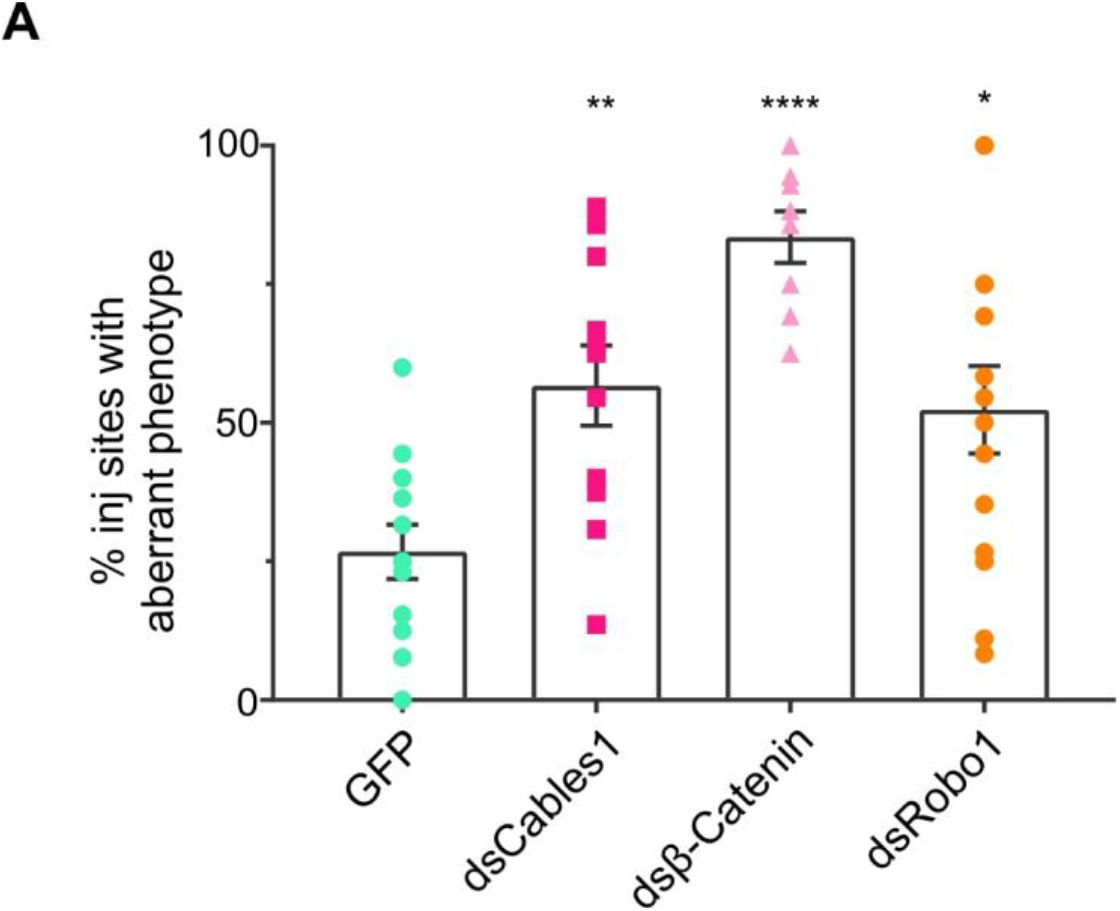
The dsRNAs derived from *Cables1, β-Catenin*, and *Robo1* efficiently perturbed axon guidance when used at regular concentration. Using the regular concentrations of 300 ng/µl of dsRNA to silence target genes effectively interfered with commissural axon guidance. Axonal trajectories were aberrant at 55.4±6.9% of the DiI injection sites after silencing *Cables1* (n=123 injection sites in N=11 embryos), 82.3±4.9% of the injection sites after silencing *β-Catenin* (n=91, N=8), and 51.1±7.7% of the DiI sites in dsRobo1-treated embryos (n=143, N=14). Embryos injected with the plasmid encoding GFP were used as controls. Pathfinding was affected only at 26.2±4.8% of the injection sites in control-treated embryos (n=125 injection sites in N=12 embryos).

**Supplementary Figure 5:**
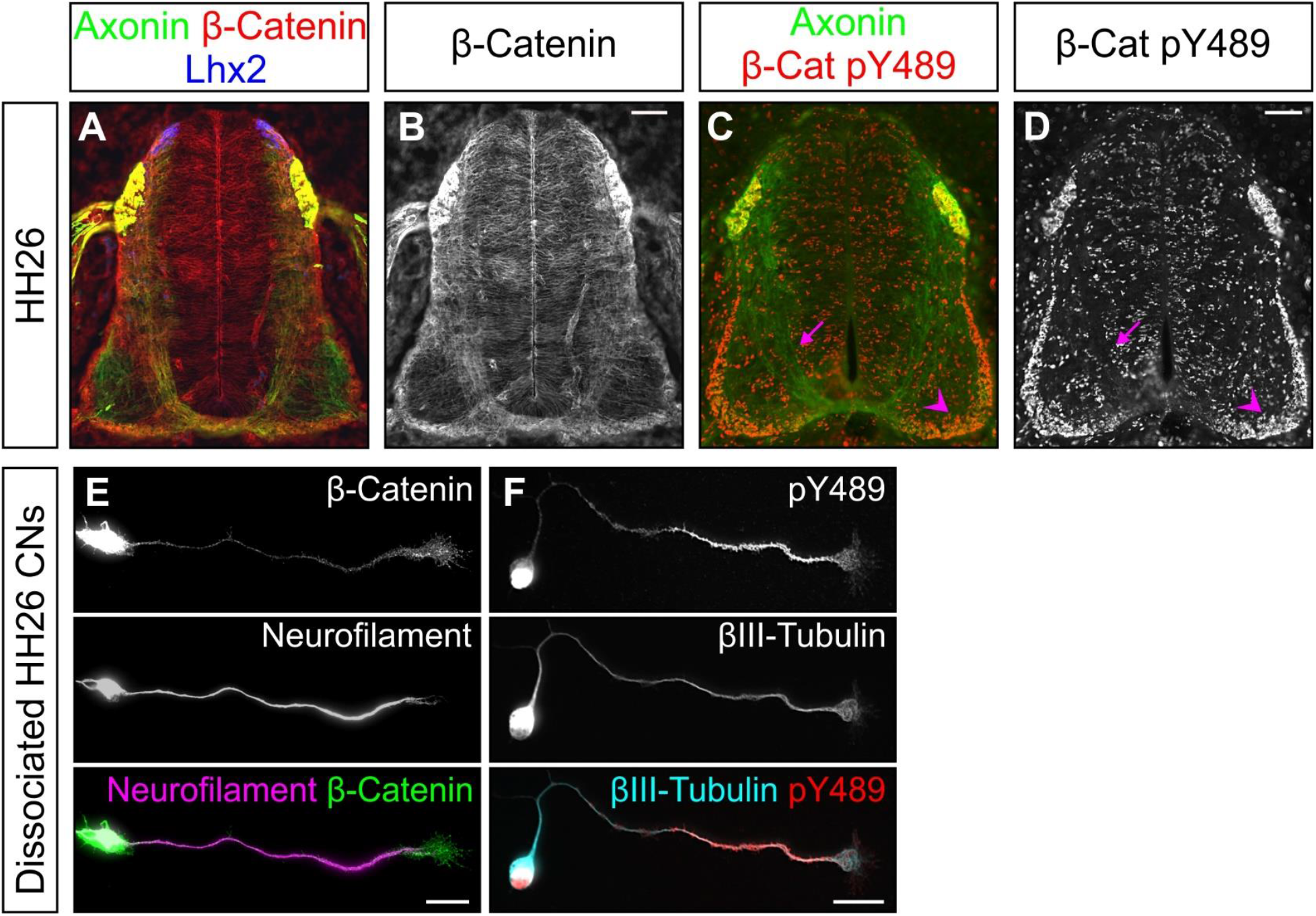
A phosphorylated form of β-Catenin, β-Catenin pY489, accumulates in distal post-crossing axons. Transverse sections of spinal cords from HH26 embryos were stained for total β-Catenin (A, B) or β-Catenin pY489 (C, D). With an antibody recognizing all forms of β-Catenin, pre- and post-crossing axons were stained (A, B). In contrast, an antibody specific for β-Catenin that is phosphorylated at Y489 revealed higher levels of -β-Catenin pY489 on post-crossing axons (arrowhead), very low levels or no β-Catenin pY489 was found on pre-crossing axons (arrows) (C, D). Commissural axons and axons from dorsal root ganglia (DRG) neurons are visualized with an anti-Axonin1 antibody (green, A, C). Staining of cultures of dissociated neurons dissected from embryos sacrificed at HH26 demonstrates the accumulation of β-Catenin pY489 in the distal axon (F), whereas levels of total β-Catenin are more homogenous along the axon (E).

**Supplementary Figure 6:**
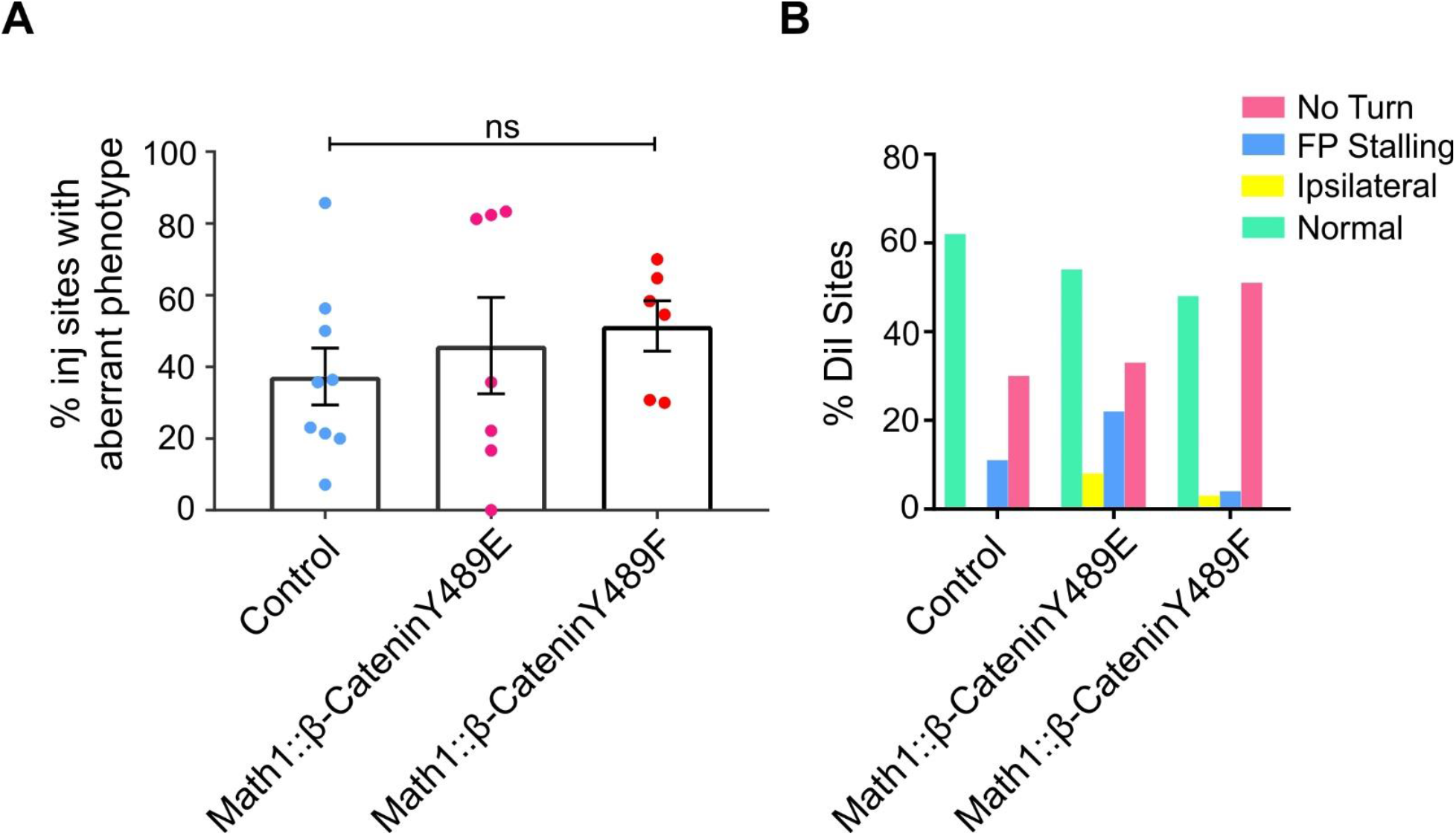
Overexpression of β-Catenin Y489 phosphomutants does not significantly affect commissural axon guidance. (A) Quantification of aberrant axon guidance phenotypes per embryo obtained after overexpression of β-Catenin Y489E (phosphomimetic) or Y489F (phosphoinhibited). None of the mutant forms of β-Catenin impaired commissural axon guidance. Data are shown as mean +/-sem. We found aberrant axonal trajectories at 37.3±7.9% of the DiI injection sites in control-treated embryos injected with the empty Math1::IRES-GFP plasmid in comparison to 45.9±13.4% in β-Catenin Y489E-expressing and 51.4±6.9% in β-Catenin Y489F-expressing embryos. ns= not significant. Tukey’s multiple comparisons test. (B) Distribution of axon guidance phenotypes. Control, N=9, n=35; β-Catenin Y489E, N=7, n=89; β-Catenin Y489F, N=6, n=73.

